# Linkage equilibrium between rare mutations

**DOI:** 10.1101/2024.03.28.587282

**Authors:** Anastasia S. Lyulina, Zhiru Liu, Benjamin H. Good

## Abstract

Recombination breaks down genetic linkage by reshuffling existing variants onto new genetic backgrounds. These dynamics are traditionally quantified by examining the correlations between alleles, and how they decay as a function of the recombination rate. However, the magnitudes of these correlations are strongly influenced by other evolutionary forces like natural selection and genetic drift, making it difficult to tease out the effects of recombination. Here we introduce a theoretical framework for analyzing an alternative family of statistics that measure the homoplasy produced by recombination. We derive analytical expressions that predict how these statistics depend on the rates of recombination and recurrent mutation, the strength of negative selection and genetic drift, and the present-day frequencies of the mutant alleles. We find that the degree of homoplasy can strongly depend on this frequency scale, which reflects the underlying timescales over which these mutations occurred. We show how these scaling properties can be used to isolate the effects of recombination, and discuss their implications for the rates of horizontal gene transfer in bacteria.

## INTRODUCTION

The statistical associations between mutations, also known as linkage disequilibrium (LD), contain a wealth of information about the evolutionary forces acting within a population (Slatkin, 2008). Chief among these is recombination, which breaks up genetic linkage by reshuffling existing variants onto new genetic backgrounds. Linkage disequilibrium has played a central role in illuminating the recombination dynamics of natural populations, from fine-scale recombination maps in sexual organisms (Chan *et al*., 2012; Coop *et al*., 2008; McVean *et al*., 2004; Myers *et al*., 2005; Spence and Song, 2019) to the rates of horizontal gene transfer in bacteria (Didelot and Falush, 2007; Didelot and Wilson, 2015; Garud *et al*., 2019; Lin and Kussell, 2017; Liu and Good, 2024; Rosen *et al*., 2015), viruses (Neher and Leitner, 2010; Romero and Feder, 2024; Turakhia *et al*., 2022; Zanini *et al*., 2015), and other microbes (Lynch *et al*., 2022; Vakhru-sheva *et al*., 2020). In addition to recombination, LD also encodes important information about the demographic history of a population (Li and Durbin, 2011; Ragsdale and Gravel, 2019; Ragsdale *et al*., 2023; Santiago *et al*., 2020) and the action of positive (Garud *et al*., 2015; Sabeti *et al*., 2002; Stephan *et al*., 2006; Wolff and Garud, 2023) or negative (Corbett-Detig *et al*., 2013; Garcia and Lohmueller, 2021; Ragsdale, 2022; Sohail *et al*., 2017) selection. However, disentangling the contributions of these forces remains challenging (Garud *et al*., 2021; Harris *et al*., 2018), since the statistical associations between mutations are only partially understood theoretically.

Much of our existing understanding of LD has focused on the pairwise correlations between alleles at different locations on the genome. These pairwise correlations are often summarized by the squared correlation coefficient,

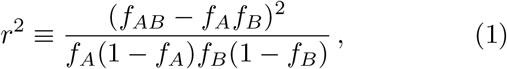

where *f*_*A*_ and *f*_*B*_ denote the marginal frequencies of the mutant alleles at each site, and *f*_*AB*_ denotes the fraction of individuals with mutant alleles at both sites (Hill and Robertson, 1968). The *r*^2^ metric and related measures like *D*′ (Lewontin, 1964) and 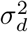 (Ohta and Kimura, 1971) quantify how the observed genomes deviate from the infinite recombination limit (also known as *linkage equilibrium*), where the alleles at each site are independently distributed across genetic backgrounds (*f*_*AB*_ ≈ *f*_*A*_ · *f*_*B*_).

The frequencies in Eq. (1) are themselves random variables that emerge from an underlying evolutionary model. Several theoretical approaches have been developed for predicting how the moments of *r*^2^ and related correlation metrics scale with the recombination rate and other parameters in particular evolutionary scenarios (Good, 2022; Lin and Kussell, 2017; Lynch *et al*., 2014; McVean, 2002; Ohta and Kimura, 1971; Ragsdale, 2022; Ragsdale and Gravel, 2019; Santiago *et al*., 2020; Song and Song, 2007; Stephan *et al*., 2006). More recent work has started to explore how these correlations vary as a function of the frequencies of the two alleles (Eberle *et al*., 2006; Good, 2022; Lynch *et al*., 2022; Rosen *et al*., 2015; Sohail *et al*., 2017; Wolff and Garud, 2023), which are increasingly accessible with the large sample sizes of modern genomic datasets (Almeida *et al*., 2021; Halldorsson *et al*., 2022; Sun *et al*., 2023). Since the frequencies of these variants are related to the time at which they arose, this frequency dependence allows us to probe how evolutionary forces contribute to LD across a range of different timescales (Good, 2022).

However, correlation metrics like *r*^2^ are just one way of summarizing the statistical associations between pairs of mutations. In principle, this information is fully contained in the two-locus haplotype frequency spectrum, *p*(*f*_*Ab*_, *f*_*aB*_, *f*_*AB*_), which is the continuous analogue of the two-locus sampling distribution that has been explored in previous work (Hudson, 2001; Ragsdale *et al*., 2018). Just as existing metrics like Tajima’s *D* and Fay and Wu’s *H* are sensitive to different portions of the singlesite frequency spectrum (Fay and Wu, 2000; Fu and Li, 1993; Tajima, 1989), other two-locus statistics will generally capture different portions of the haplotype frequency spectrum (Good, 2022; Ragsdale and Gravel, 2019), and may therefore be useful for teasing out the contributions of different evolutionary forces.

For example, another class of summary statistics derives from the four-gamete test (Hey and Wakeley, 1997; Hudson and Kaplan, 1985; Neher and Leitner, 2010; Vakhrusheva *et al*., 2020), which asks whether all four combinations of alleles are present within a sample. This functions as test for homoplasy, since the fourth combination can only be produced via recombination or recurrent mutations. While the four-gamete test is usually viewed as a binary readout, we can also define a more graduated version,

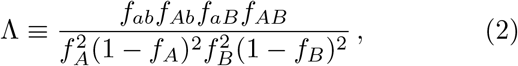

which is a continuous function of the four haplotype frequencies. Like the original four gamete test, this Λ statistic vanishes in the absence of recombination or recurrent mutation, but is normalized so that it approaches one under linkage equilibrium (*f*_*AB*_ ≈ *f*_*A*_ · *f*_*B*_). In this way, Eq. (2) quantifies the deviation from the zero recombination limit, similar to how *r*^2^ captures the deviation from the infinite recombination limit. This suggests that homoplasy metrics like Λ could be particularly useful for isolating the effects of recombination. The degree of homoplasy is also important in other evolutionary contexts: it determines the “softness” of selective sweeps from standing genetic variation (Hermisson and Pennings, 2017), and can reveal the presence of genetic incompatibilities between loosely linked loci (Corbett-Detig *et al*., 2013).

Despite these attractive properties, our quantitative understanding of homoplasy statistics like Eq. (2) remains limited, even in the simplest evolutionary scenarios. While some properties of the four gamete test can be derived using coalescent theory (Hey and Wakeley, 1997; Myers and Griffiths, 2003), it is difficult to extend these calculations to larger sample sizes, or to account for natural selection or recurrent mutation. Our limited understanding of these effects leaves many basic questions unresolved: How does the buildup of homoplasy compare with the decay of LD as the distance between sites increases? Does negative selection change this picture? Can we distinguish recombination from recurrent mutation using quantitative metrics like Eq. (2)? And finally, how do the answers to these questions depend on the frequencies of the two alleles?

Here, we address these questions by generalizing a recently developed framework for modeling frequency-resolved LD (Good, 2022) to study homoplasy statistics like Eq. (2). We focus on weighted moments of Λ, where the weights are chosen to single out particular frequency scales of the underlying alleles. Using this approach, we derive analytical expressions that predict how these homoplasy statistics depend on the rates of recombination and recurrent mutation, and the additive and epistatic fitness costs of the mutations. We show how these approaches can be generalized to predict the full distribution of Λ, conditioned on the marginal frequencies of the two alleles. We conclude by discussing the implications of these results for measuring recombination dynamics in large microbial datasets.

### MODEL AND ANALYSIS

We investigate the dynamics of homoplasy statistics like Eq. (2) in a two-locus Wright-Fisher model under the joint action of mutation, recombination, negative selection, and genetic drift. We consider a panmictic population of *N* haploid individuals with two biallelic loci, *a/A* and *b/B*, that each acquire mutations at rate *μ* per individual per generation. We will restrict our attention to cases where *Nμ* ≪ 1, which ensures that the pairwise heterozygosity at each site is also low (Ewens, 2004). We assume that the *A* and *B* alleles lower the fitness of an individual by *s*_*A*_ and *s*_*B*_, respectively, while mutations at both loci impose a total cost *s*_*AB*_ = *s*_*A*_ + *s*_*B*_ + *ϵ*_*AB*_, with *ϵ*_*AB*_ denoting the amount of epistasis. Finally, we assume that the two loci recombine at a total rate *R* per genome per generation, which depends on the coordinate distance *ℓ* between the two loci. Most of our results will be independent of the functional form of *R*(*ℓ*), provided that we write our expressions in terms of the map distance *R*. These assumptions yield a standard two-locus Wright-Fisher model (Appendix A) whose equilibrium distribution we will denote by *p*(*f*_*Ab*_, *f*_*aB*_, *f*_*AB*_).

To explore how homoplasy emerges across a range of different timescales, we extend the approach introduced in Good (2022) and consider weighted moments of Λ that condition on the marginal frequencies of the alleles at each of the two loci. We consider two different classes of weighting functions in this work. The first class, which was previously introduced in Good (2022), allows us to focus on the dynamics when the minor alleles at both sites are rare. The weighting function in this case is defined as

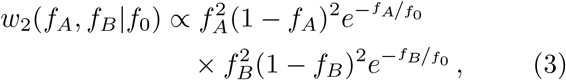

where *f*_0_ is a characteristic allele frequency scale, and the proportionality constant is chosen such that the expectation of *w*_2_(*f*_*A*_, *f*_*B*_| *f*_0_) under the equilibrium distribution *p*(*f*_*Ab*_, *f*_*aB*_, *f*_*AB*_) is normalized to one. The average value of Λ under this weighting scheme is therefore given by

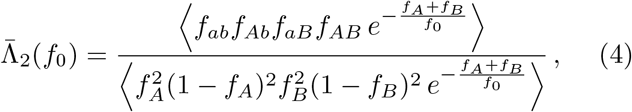

where the angle brackets ⟨·⟩ denote the expectation under the equilibrium distribution *p*(*f*_*Ab*_, *f*_*aB*_, *f*_*AB*_). The exponential weighting terms in Eq. (4) act like a soft step function, preferentially excluding alleles with frequencies ≳ *f*_0_. The exponential cutoff has convenient analytical properties that we will exploit below, but many of our qualitative results will apply for other choices of the cutoff function provided that they remain sufficiently sharp.

In addition to Eq. (4), we also consider a second class of weighting functions that allow us to condition on cases where only one of the two alleles (e.g. *A*) is rare, while the other is at an intermediate frequency. The weighting function in this case is defined as

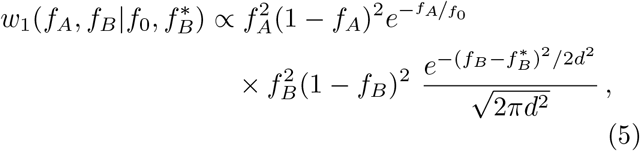

where *f*_0_ and 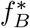 are a pair of allele frequency scales satisfying 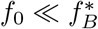, and *d* is a characteristic width that determines the range of *f*_*B*_ values that contribute to *w*_1_. We focus on small values of *d* such that the Gaussian term in Eq. (5) approaches a Dirac delta function, which forces 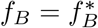. The average value of Λ under this “single-rare” weighting scheme is then given by

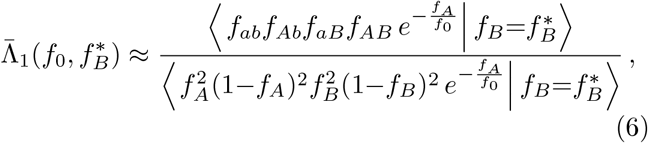

where the angle brackets again denote an average over the equilibrium distribution *p*(*f*_*Ab*_, *f*_*aB*_, *f*_*AB*_).

The weighted average in Eq. (4) can be straightforwardly found from the moment generating function,

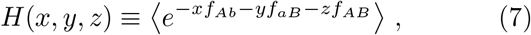

using the identity

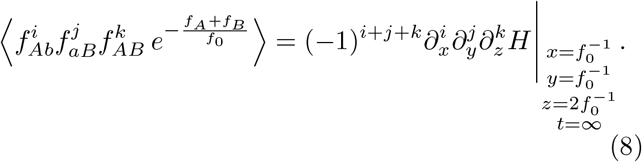

The conditional averages in Eq. (6) obey a similar relation involving the conditional generating function,

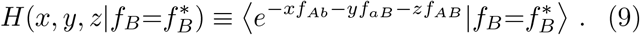

We calculate these quantities by extending the analytical approach employed in Good (2022), which yields a perturbative solution of Eq. (8) that applies in the limit that the frequencies of the alleles are rare (*f*_0_ ≪ 1). The evolutionary dynamics greatly simplify in this limit because only a few distinct classes of frequency trajectories will end up contributing to the averages in Eq. (8) (Fig. 1), each of which can be associated with a corresponding term in the perturbation expansion of *H*(*x, y, z*). We derive these formal solutions in Appendices B and F, respectively. In the following sections, we use these results to develop predictions for Λ in different evolutionary scenarios.

**FIG. 1.**
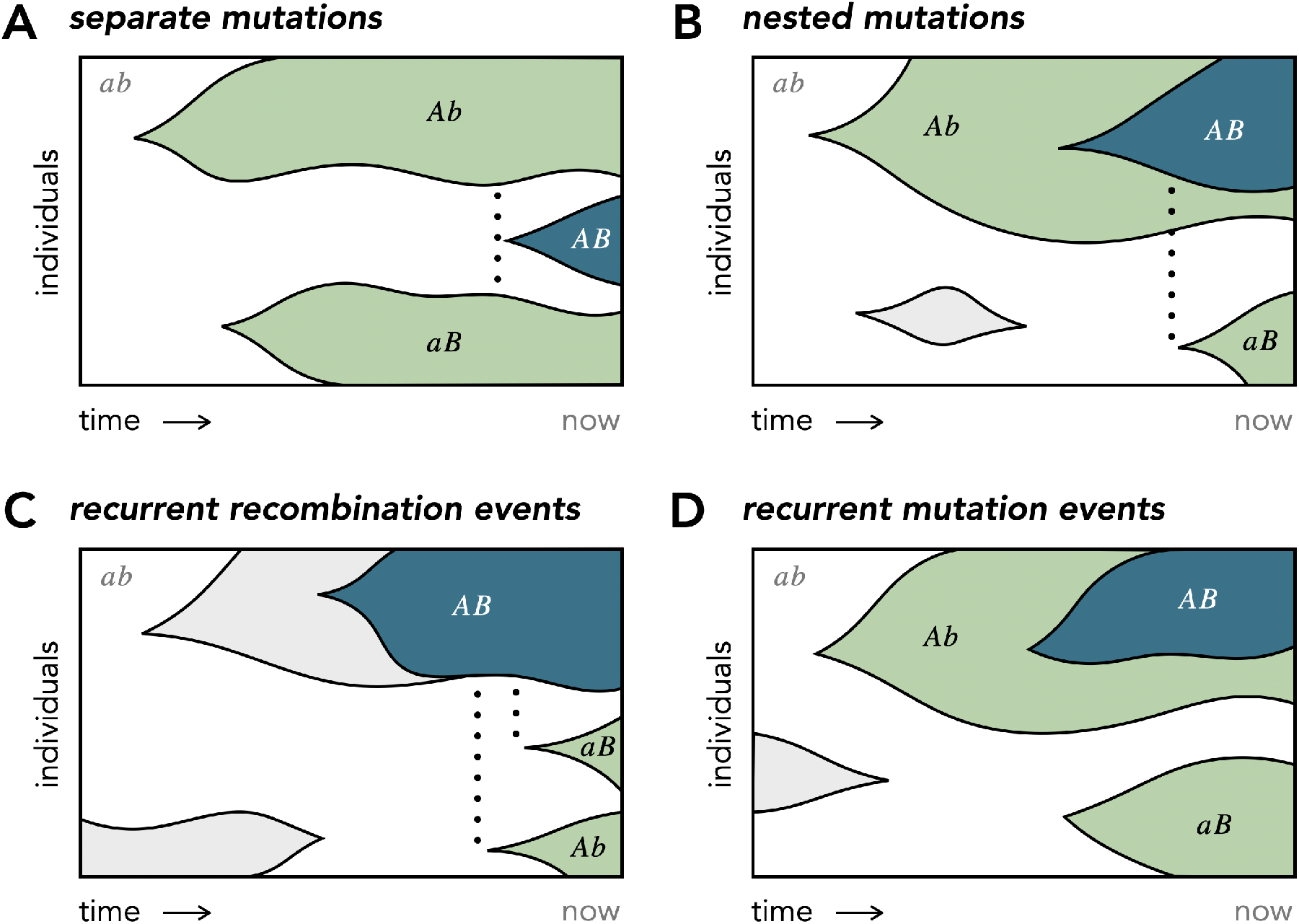
Schematic of lineage dynamics that contribute to homoplasy when mutant alleles are rare. **(A)** separate mutations. *A* and *B* mutations arise separately on the wildtype background *ab*. Before going extinct, they recombine and produce the double-mutant lineage *AB* (vertical dotted line). **(B)** nested mutations. The wildtype population first acquires mutation *A* and then *B* mutation occurs on the *Ab* background. The double mutant then recombines with the wildtype and generates the missing single-mutant haplotype *aB*. All three mutant lineages are still segregating at the time of observation. **(C)** recurrent recombination events. The wildtype population acquires mutation *A* and then *B* mutation occurs on the *Ab* background. The double mutant then recombines with the wildtype and produces *aB*, however, by that time the *Ab* lineage has gone extinct. The presence of all four haplotypes therefore necessitates an additional recombination event. An analogous diagram exists for the case of separate mutations with two recombination events (not shown). **(D)** recurrent mutation events. Mutation *B* arises twice on different backgrounds, *ab* and *Ab*, to produce the fourth haplotype.

## RESULTS

### Neutral alleles

The simplest behavior occurs in the absence of selection (*s*_*A*_, *s*_*B*_, *s*_*AB*_ = 0), when recurrent mutations can be neglected (*Nμ* → 0). In this case, 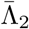 in Eq. (4) will only depend on the population-scaled recombination rate *NR* in addition to the frequency scale *f*_0_. When *f*_0_ ≲ 10%, we find that the solution for 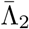 collapses onto a singleparameter curve,

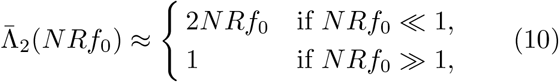

which transitions from a *recombination-limited* regime 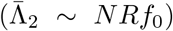 when *NRf*_0_ ≪ 1 to a *recombination-dominated* regime 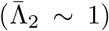 when *NRf*_0_ ≫ 1 (Fig. 2; Appendices C and D). The transition between these two limits occurs when *NRf*_0_ ∼ 𝒪(1), where the full numerical solution is necessary to obtain quantitative agreement with simulations. Since 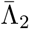 only depends on the compound parameter *NRf*_0_, these results imply that a lower frequency scale *f*_0_ can mimic the effects of a lower recombination rate, and vice versa. In particular, a pair of sites can be in the recombination-limited regime even if their nominal recombination rate is high (*NR* ≫ 1), provided that the frequency scale is sufficiently low (*f*_0_ ≲ 1*/NR*).

**FIG. 2.**
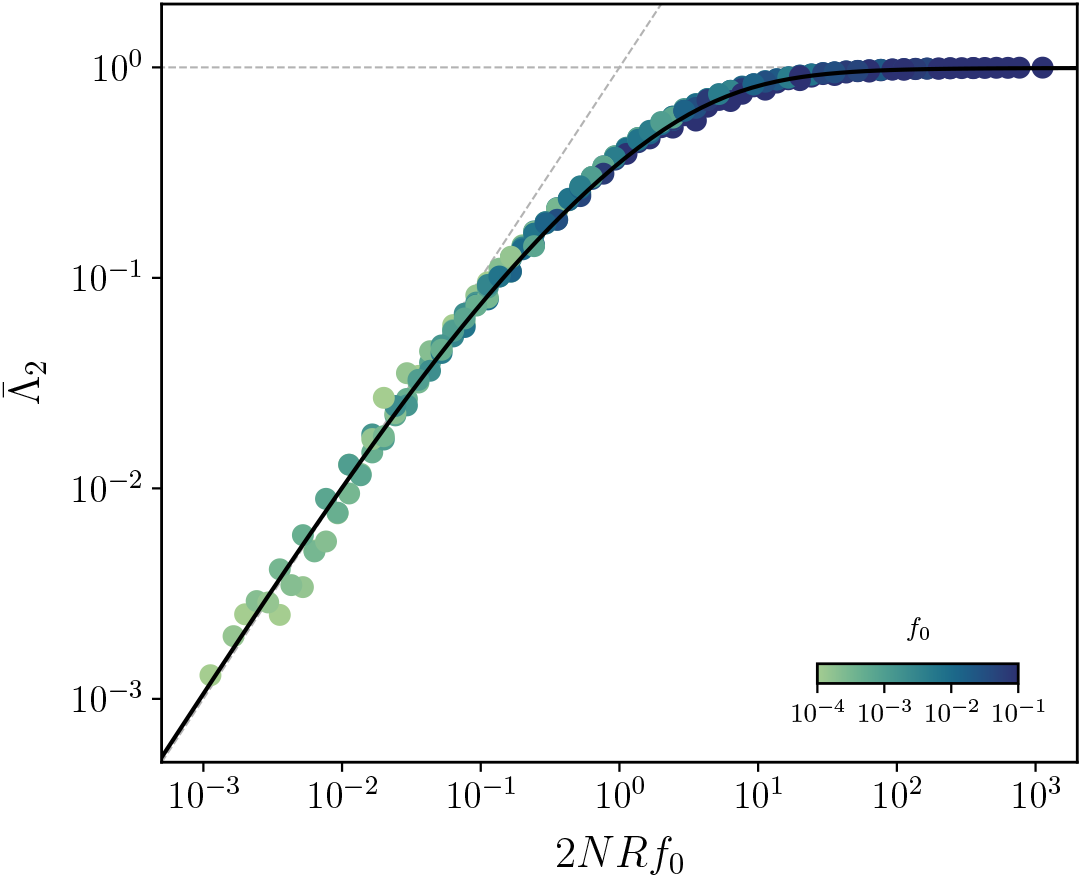
Frequency-resolved homoplasy, 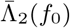, for pairs of neutral alleles in the infinite sites limit. Points denote the results of forward-time simulations (Appendix A) with *N* = 10^6^ and different combinations of *R* and *f*_0_; each point represents an average over 10^9^ pairs of loci. The fact that different combinations of parameters collapse onto a single curve suggests that 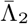 is primarily determined by the compound parameter *NRf*_0_. The solid black line shows the theoretical prediction from Eq. (C6), while the dashed gray lines show the asymptotics from Eq. (10).

We can develop an intuition for the behavior in Eq. (10) by considering the haplotype frequency dynamics that contribute to 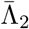. When recombination is rare, 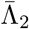 will be dominated by trajectories involving a single recombination event. In the infinite sites limit (*Nμ* → 0), there are only two distinct ways to produce all four haplotypes in the population. In the first case, separate mutations on the wildtype background can create a pair of single-mutant lineages, *Ab* and *aB*, which recombine with each other to produce the double-mutant haplotype *AB* (Fig. 1A). Alternatively, the double-mutant haplotype could be produced by a nested mutation within one of the single-mutant backgrounds, which then recombines with the wildtype to create the missing fourth haplotype (Fig. 1B).

We can estimate the contributions of each scenario to homoplasy statistics like 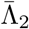 using the heuristic approach described in Good (2022). The averages in the numerator and denominator of Eq. (4) can be estimated by multiplying the probability of each event by the typical haplotype frequencies it is associated with. In both cases, the averages will be dominated by mutations that arose within the last ∼*Nf*_0_ generations and drifted to a characteristic frequency scale ∼ *f*_0_ (Good, 2022). In the separate mutations case (Fig. 1A), the two single mutants each arise at rate ∼ *Nμ*, while the recombinant double mutants are produced at rate 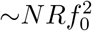. In the nested mutations case (Fig. 1B), the second mutation is produced at rate ∼*Nμf*_0_, while the remaining recombinant haplotype is produced at rate ∼ *NRf*_0_ · 1 when the double mutant recombines with the wildtype. The higher production of recombinants in this case is exactly balanced by the lower rate of producing nested mutations, resulting in similar overall contributions to the numerator of Eq. (4):

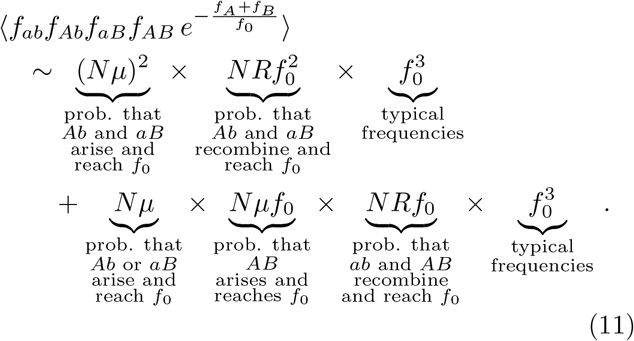

A similar calculation shows that the denominator of Eq. (4) is given by

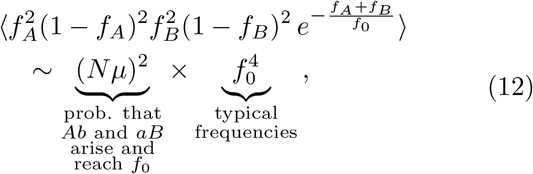

so that the ratio between the two expressions yields the 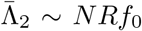 dependence observed in Eq. (10) when *NRf*_0_ ≪ 1.

The strong recombination regime can be understood using a similar approach, except that we now have to account for the greater loss of *AB* individuals due to recombination. This outflow imposes an effective fitness cost *R* on the double mutant, which prevents it from rising above a frequency ∼1*/NR* (Good, 2022). When this maximum frequency is less than ∼ *f*_0_, the numerator in Eq. (11) must instead be replaced by

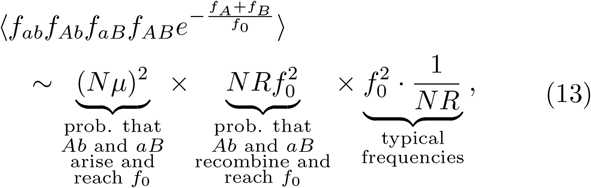

which divided by Eq. (12) yields the 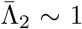 scaling observed in Eq. (10) when *NRf*_0_ ≫ 1.

As the rate of recombination becomes even larger 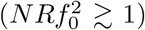, multiple double-mutant lineages will start to be produced by recombination every generation. In this case, the total size of the *AB* haplotype will be determined by a balance between the production rate of new recombinants 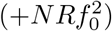 and their loss due to further recombination with the wildtype (− *NRf*_*AB*_). The balance between these terms occurs when 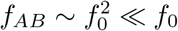, which is equivalent to the condition that the *A* and *B* mutations are in *quasi-linkage equilibrium* (Good, 2022). In this case, our normalization convention in Eq. (2) ensures that Λ is close to one, so that the average 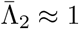 as well. Since this average value is the same as in the *NRf*_0_ ≫ 1 case, higher moments of Λ are required to observe the transition to the quasi-linkage equilibrium regime. We consider this case in more detail in a separate section below.

### Incorporating negative selection

We are now in a position to understand how negative selection on the mutants changes the behavior observed above. We begin by considering the simplest case, where the *A* and *B* mutations have the same fitness cost (*s*_*A*_ = *s*_*B*_ = *s*) and there is no additional epistasis (*s*_*AB*_ = 2*s*). In this case, we find that the solution for 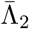 exhibits a similar transition from a linear dependence on *R* when *R* → 0 to a saturated regime when *R*→ ∞ (Fig. 3). However, the location of this transition now depends on the relative strengths of negative selection and genetic drift. When *Nsf*_0_ ≪ 1, selection will not have had a chance to alter the frequencies of the mutations while they were drifting to their present-day frequencies (∼ *f*_0_). This implies that 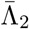 will remain close to the neutral result in Eq. (10) (Fig. 3, left). Since the boundary of this regime depends on the compound parameter *Nsf*_0_, even strongly deleterious mutations (*Ns* ≫ 1) can behave effectively neutrally if *f*_0_ is chosen to be sufficiently low.

**FIG. 3.**
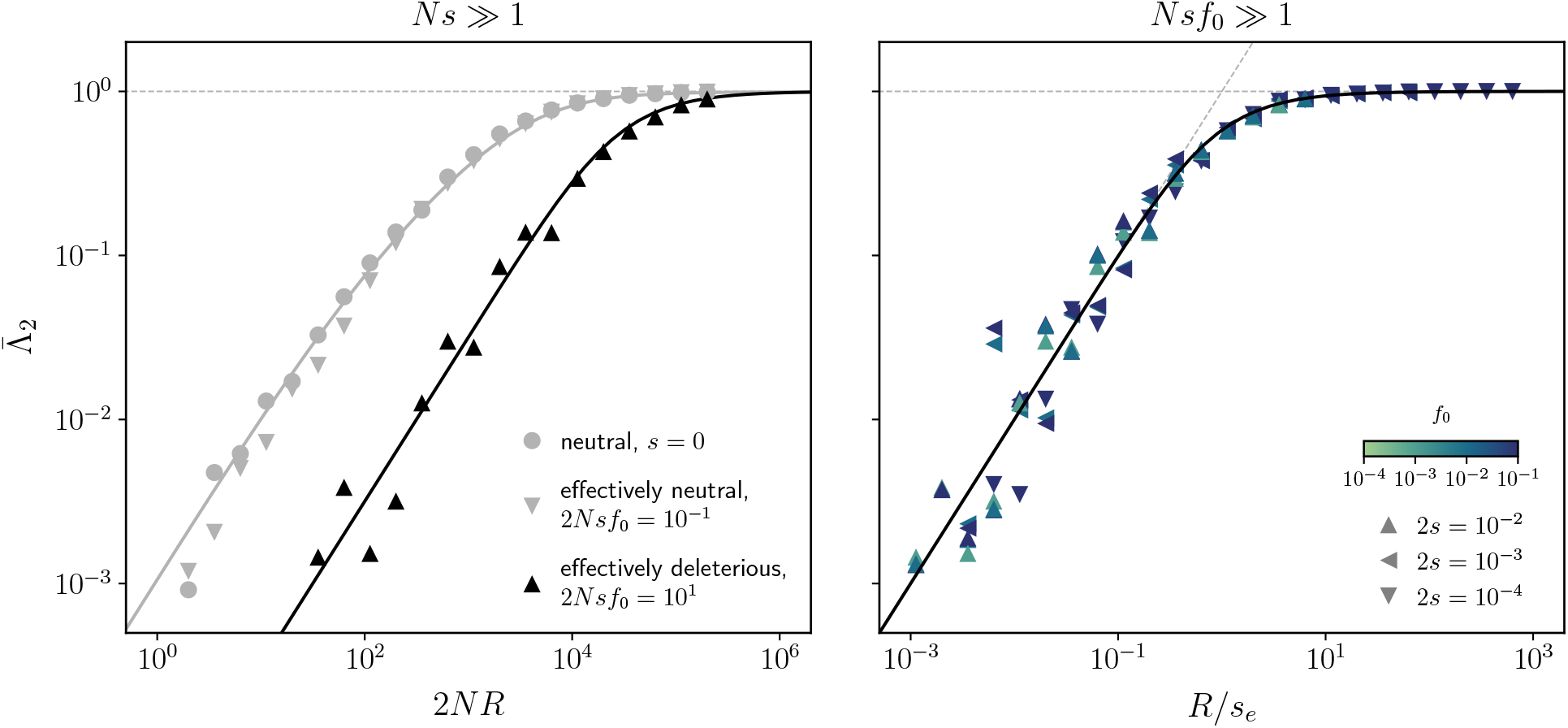
Frequency-resolved homoplasy, 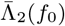, for pairs of negatively selected alleles. **Left:** 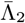 as a function of the population-scaled recombination rate (*NR*) when *s*_*A*_ = *s*_*B*_ = *s, s*_*AB*_ = 2*s*. Symbols denote the results of forward-time simulations with *N* = 10^6^, *f*_0_ = 10^−3^, and different values of *s*. Solid lines show the theoretical predictions for the strong selection (*Nsf*_0_ ≫ 1; black, Eq. E14) and weak selection (*Nsf*_0_ ≪ 1; grey, Eq. C6) limits. The fact that the grey symbols collapse onto the same curve illustrates that even strongly deleterious mutations (grey triangles, *Ns* ∼ 10^2^) can behave effectively neutrally if *Nsf*_0_ ≪ 1. **Right:** an analogous version of the left panel showing 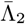 as a function of the selection-scaled recombination rate, *R/s*_*e*_, with *s*_*e*_ ≡ (60*/*19)*s*. Symbols denote the results of forward-time simulations with *N* = 10^6^ and different combinations of *R, s*, and *f*_0_. Similar to Fig. 2, the fact that different combinations of parameters collapse onto a single curve suggests that 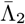 is primarily determined by the compound parameter *R/s*_*e*_ in the strong selection regime (*Nsf*_0_ ≫ 1). The solid line shows the theoretical prediction from Eq. (E14), while the grey dashed lines show the asymptotics from Eq. (14).

In the opposite case, when selection is strong compared to drift (*Nsf*_0_ ≫ 1), we find that 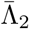 can be expressed as

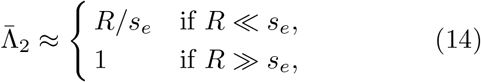

where *s*_*e*_ ≡ (60*/*19)*s* is an effective fitness cost (Fig. 3; Appendix E). The functional form of this expression suggests that negative selection has a similar effect as imposing a frequency threshold at 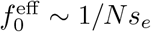. The primary difference occurs in the narrow crossover region where 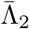 approaches saturation: a comparison of the two curves shows that the transition between the recombinationlimited 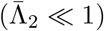 and recombination-dominated 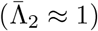 regimes is slightly sharper in the presence of strong negative selection (Fig. 3, left).

Interestingly, we find that in many other cases of strong selection, individual fitness costs can be absorbed by the effective cost *s*_*e*_, so that the asymptotic behavior of 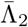 is still well approximated by the limits in Eq. (14) (Appendix E). This continues to hold for a range of epistatic interactions, as long as single mutations are not much more deleterious alone than in combination (*s*_*A*_, *s*_*B*_ ≲ *s*_*AB*_).

The simplicity of this behavior can be understood by revisiting our heuristic picture above. The frequency trajectories of deleterious alleles are similar to those of neutral alleles, except that negative selection prevents them from growing to frequencies much larger than the drift barrier at ∼ 1*/Ns* (Fisher, 2007). When this maximum frequency is smaller than *f*_0_, the sizes of the single-mutant lineages will be capped at 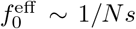 instead of the nominal threshold at *f*_0_. Similarly, the typical frequency of the double mutant will depend on the relative strengths of selection and recombination,

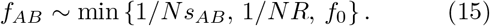

Substituting these typical frequencies into Eqs. (11) and (12) yields the asymptotic behavior in Eq. (14).

We note, however, that while this simple expression captures the behavior of 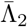 across a wide range of parameter space, more complex scenarios are possible. One notable exception occurs for strong antagonistic epistasis, where the double mutant is much less costly than either of the single mutants alone (*s*_*AB*_ ≪ *s*_*A*_, *s*_*B*_). In the extreme case where the single mutants are strongly deleterious (*Nsf*_0_ ≫ 1) but the double mutant is effectively neutral (*Ns*_*AB*_*f*_0_ ≪ 1), the population must first cross a “fitness valley” (Weissman *et al*., 2009, 2010) to generate the double-mutant haplotype. Once this lucky double mutant arises, it can drift to much higher frequencies than the single-mutant lineages, which are likely to go extinct by the time that the double mutant is eventually sampled. In order for all four haplotypes to be present in the population, the surviving double mutant will have to recombine with the wildtype population closer to the time of sampling to regenerate the single-mutant lineages (Fig. 1C). These extra recombination events lead to a faster-than-linear dependence on *R* in the recombination-limited regime (*R* ≪ *s*, Fig. 4). Moreover, while 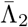 still saturates at one when *R*→ ∞, we find that it can exceed this value in the intermediate region where *R* ∼ *s* (Fig. 4).

**FIG. 4.**
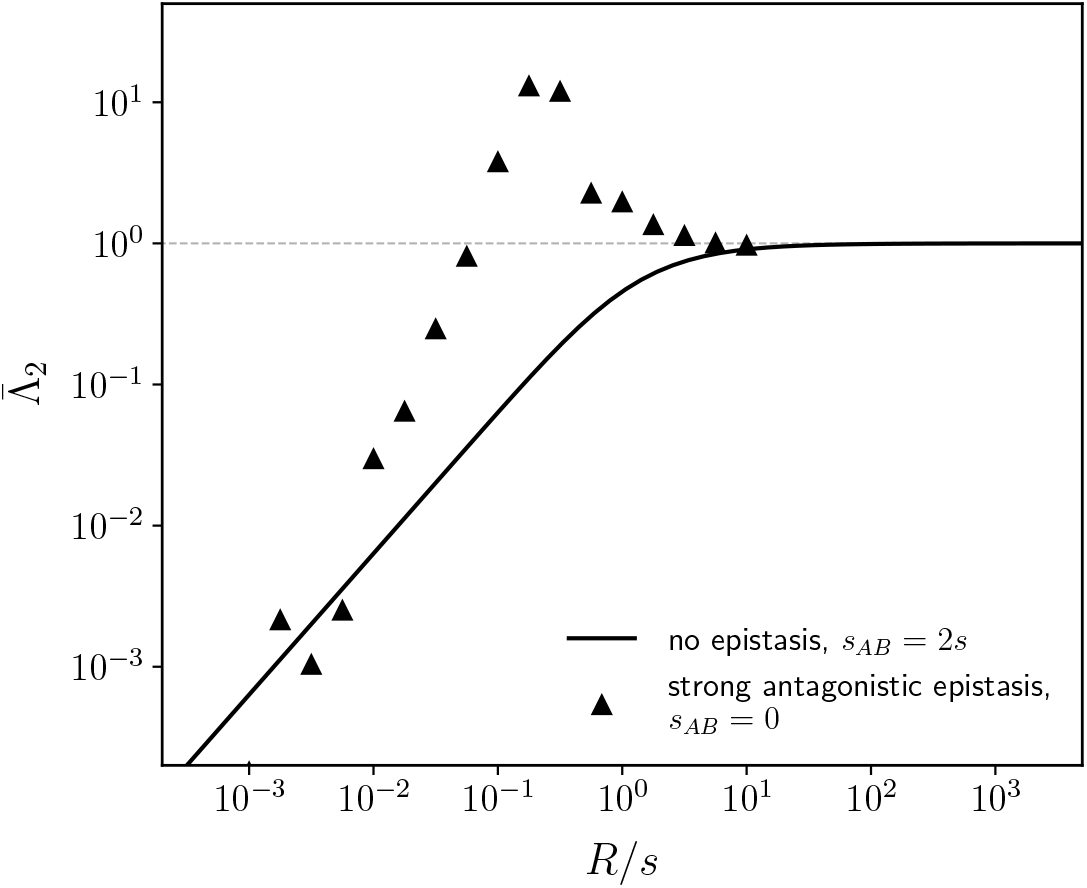
Homoplasy under strong antagonistic epistasis. An analogous version of Fig. 3, right panel for *s*_*A*_ = *s*_*B*_ = *s* = 10^−2^, *s*_*AB*_ = 0, and *N* = 10^6^. Symbols denote the results of forward-time simulations across a range of recombination rates *R* and *f*_0_ = 10^−2^. For comparison, the solid line shows the theoretical prediction from Eq. (E14) for the additive case in Fig. 3 (*s*_*AB*_ = 2*s*). Strong antagonistic epistasis changes the functional form of 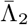 compared to the additive case: at small recombination rates, 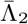 grows faster than linearly with *R*, and can temporarily exceed one (dashed line) before returning to the recombination-dominated limit when *R ≫ s*.

### Effects of recurrent mutations

Our analysis has so far focused on the infinite sites limit (*Nμ*→ 0) where the recombination was the only way to generate all four haplotypes in the population (Fig. 1A-C). However, at small but finite values of *Nμ*, recurrent mutations at either *A* or *B* locus can also create the fourth haplotype (Fig. 1D). This poses challenges for interpreting homoplasy statistics like Λ, since recurrent mutations can obscure signals of recombination, and vice versa. In this section we extend our heuristic approach to account for these effects, and show how the scaling of 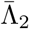 can help us distinguish between these otherwise confounding processes.

Recall that we can estimate the averages in 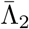 by calculating the probability that all four haplotypes arise in a population and multiplying it by their typical frequencies. At small mutation rates (*Nμ* ≪ 1), recurrent mutations will not affect the typical haplotype frequencies, but they will still alter the rate at which these haplotypes are produced. In order to produce all four combinations of alleles, at least three mutation events must happen: the wildtype population must generate a mutation at both sites and one of the single mutants must acquire an additional nested mutation. When all of these mutations are neutral, their contribution to the numerator of 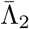 is given by a generalization of Eq. (11),

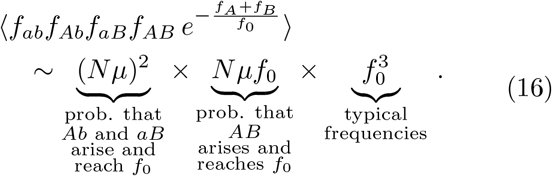

Dividing this result by the denominator in Eq. (12), we find that recurrent mutations cause 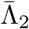 to saturate at a lower limit of ∼*Nμ*. This signal will overwhelm the contribution from recombination when *Nμ* ≫ *NRf*_0_, which leads to a modified version of Eq. (10),

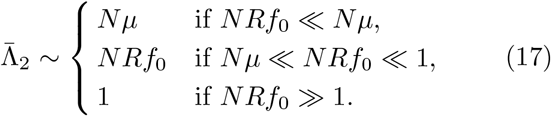

This result shows that recombination can be distinguished from recurrent mutation by examining the scaling behavior of 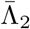. At small values of *NRf*_0_, recombination leads to a linear dependence on *f*_0_ and *R*, while recurrent mutation yields a constant value. Moreover, since *Nμ* is small, recurrent mutation will not affect the crossover to the saturated regime when *NRf*_0_ ∼ 1 (Fig. 5, left panel). Similar results apply for strong negative selection, with recurrent mutations having a negligible effect once 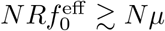 (Fig. 5, right panel). These differences in scaling arise from the graduated nature of the Λ statistic in Eq. (2), which weights large amounts of homoplasy more strongly than the small amounts produced by recurrent mutation. This suggests that the scaling behavior of 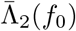 could provide a more robust signal of recombination than binary measures like the four-gamete test.

**FIG. 5.**
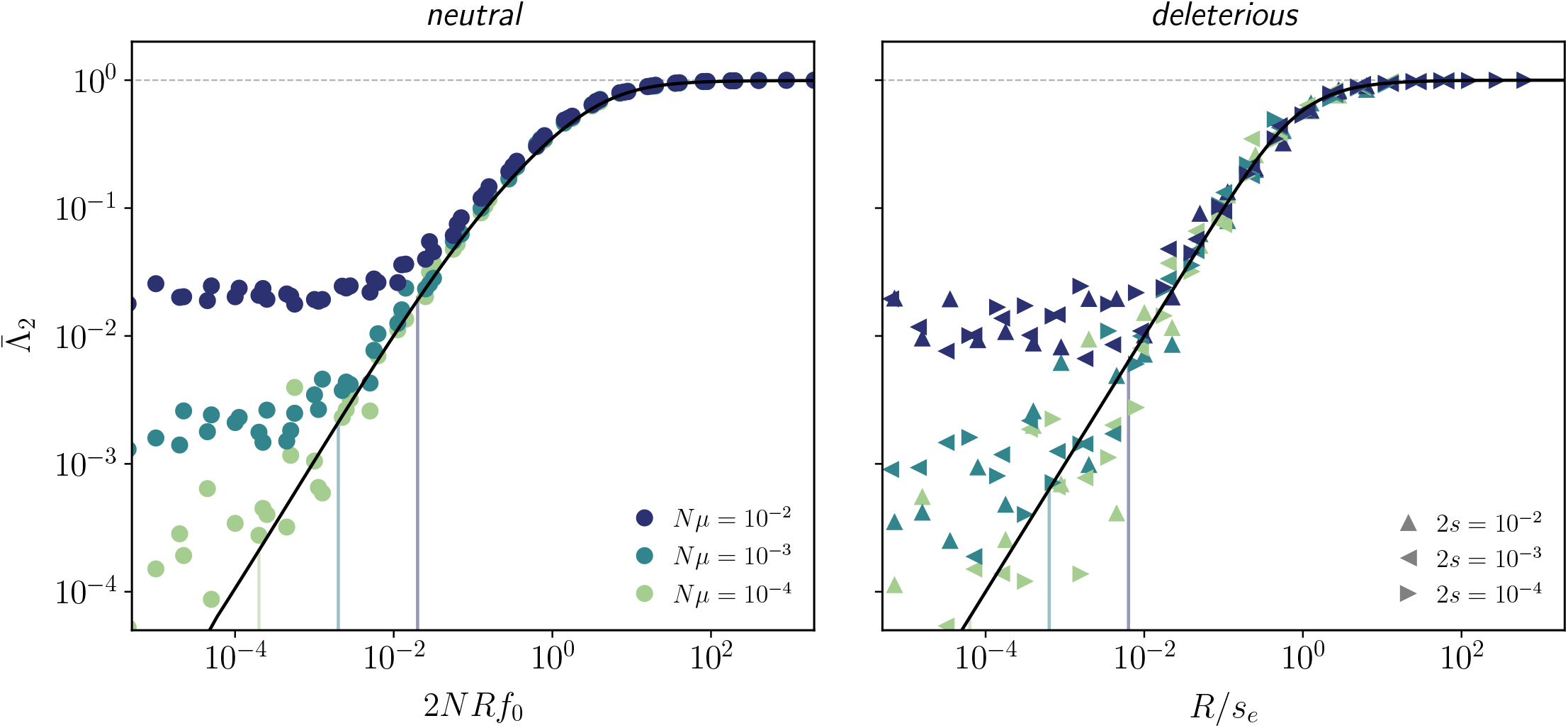
Frequency-resolved homoplasy, 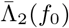, in the presence of recurrent mutations. **Left:** Analogous version of Fig. 2 for *Nμ* ∈ {10^−4^, 10^−3^, 10^−2^}. Symbols denote the results of forward-time simulations of neutral alleles for *N* = 10^6^ and *f*_0_ ∈ {10^−4^, 10^−3^, 10^−2^}. The black line shows the infinite sites prediction from Eq. (C6), while the colored lines indicate the corresponding positions where the infinite sites theory is predicted to break down (*NRf*_0_ ∼ *Nμ*). When *NRf*_0_ ≪ *Nμ*, recurrent mutations provide the dominant contribution to homoplasy, and 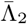 approaches a constant value ∼*Nμ*. However, as long *Nμ* ≪ 1, recurrent mutations do not affect the value of 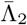 at higher values of *NRf*_0_, including the transition to the saturated regime when *NRf*_0_ ∼ 1. **Right:** an analogous version of the left panel for additive strongly deleterious alleles (*s*_*A*_ = *s*_*B*_ = 2, *s*_*AB*_ = 2*s*). Symbols denote the results of forward-time simulations for *N* = 10^6^ and *f*_0_ ∈ {10^−4^, 10^−3^, 10^−2^} across a range of recombination rates *R* and selection coefficients *s*; colors are the same as in the left panel. The black line shows the infinite sites prediction from Eq. (E14), while the colored lines indicate the analogous positions where the infinite sites theory is predicted to break down (*R/s*_*e*_ ∼ *Nμ*). These results suggest that recombination can be distinguished from recurrent mutation using the scaling behavior of 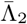.

### Distribution of Λ and the transition to linkage equilibrium

While the average in Eq. (4) contains significant information about the haplotype dynamics within a population, the full distribution of Λ can provide additional insight into the evolutionary forces at play. In this section, we explore the distribution of Λ conditioned on both alleles being rare (*f*_*A*_, *f*_*B*_ ≪ 1). We will constrain our analysis to neutral dynamics in the infinite sites limit, although it can be extended to account for certain forms of negative selection.

When recombination is frequent (*NR* ≫ 1*/f*_*A*_, 1*/f*_*B*_), double-mutant lineages are typically short-lived compared to the single-mutant lineages that produce them. In this case, previous work has shown that the total frequency of the double mutant approaches a local equilibrium,

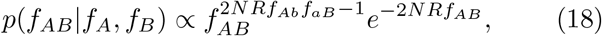

that depends on the current values of *f*_*A*_ and *f*_*B*_ (Good, 2022). When *f*_*A*_ = *f*_*B*_ = *f*_0_, the conditional distribution of Λ will therefore follow a Gamma distribution,

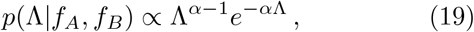

with shape parameter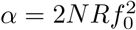.

The average of Eq. (19) is always equal to one, consistent with our previous result for recombination-dominated regime in Eq. (10). However, Eq. (19) implies that the distribution of Λ transitions between two qualitatively distinct regimes depending on the shape parameter *α* (Fig. 6A). When 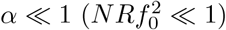, the distribution of Λ contains a large peak near zero, with an exponential cutoff at 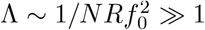. The probability mass near zero corresponds to scenarios where only three haplotypes are present at appreciable frequencies, while the exponential tail reflects the size distribution of a single *AB* lineage. While the realized values of Λ can be much larger than one in this regime, the smaller probability of these events brings the average value of 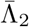 back down to one.

**FIG. 6.**
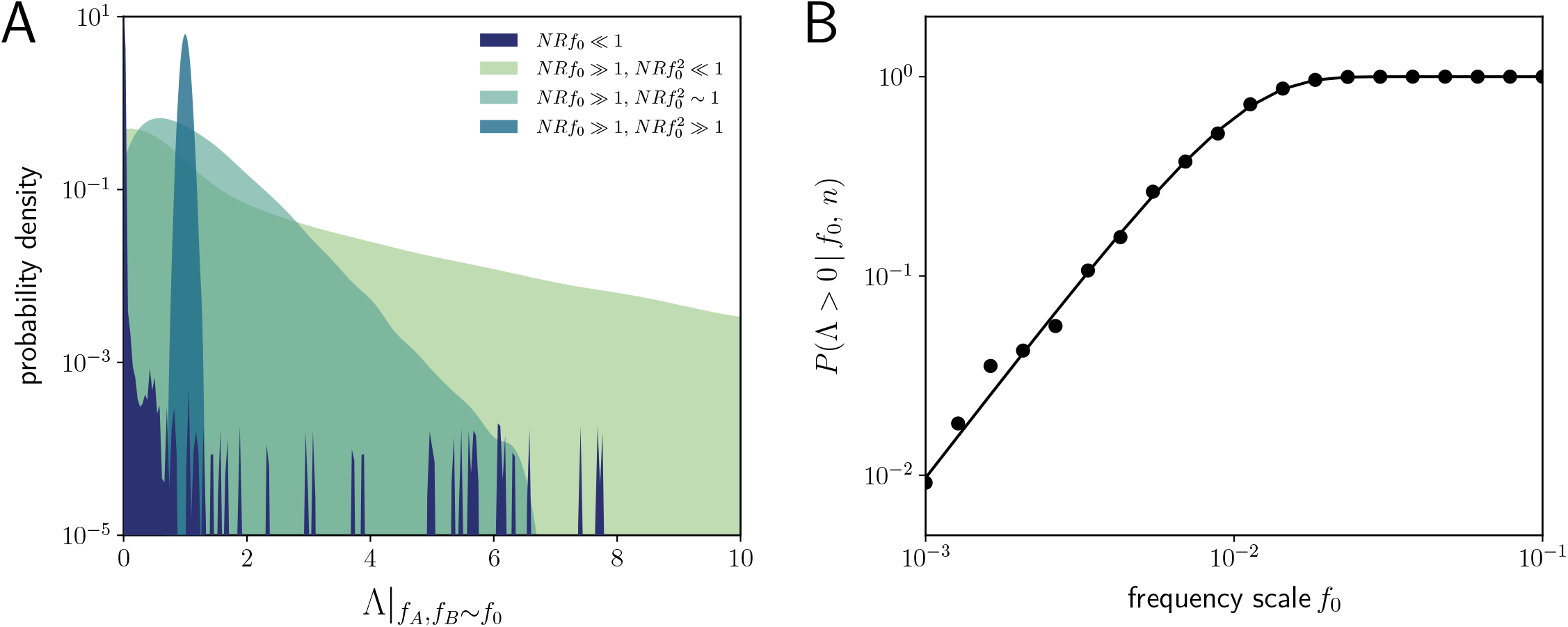
Conditional distribution of Λ when both alleles are rare. **(A)** Kernel density estimates of the distribution of Λ from simulations, conditioned on both alleles falling in a narrow range of frequencies near *f*_0_. The probability mass at Λ = 0 reflects the fraction of simulation runs where only three haplotypes were observed. Simulations were performed for neutral mutations with *N* = 10^6^, *R*≈ 5.6 · 10^−7^, *f*_0_ ≈ 1.5 · 10^−2^ (purple) and *N* = 10^6^, *R* ≈ 5.6 · 10^−2^, *f*_0_ ≈1.5 · 10^−3^ (light green), *f*_0_ ≈ 4.7 · 10^−3^ (dark green), *f*_0_ ≈ 4.7 · 10^−2^ (teal). When recombination is rare (*NRf*_0_ ≪ 1, purple), the distribution of Λ has a long tail that reflects the size of the occasional double-mutant lineage (*f*_*AB*_ ≲ *f*_0_). As the rate of recombination becomes larger (*NRf*_0_ ≳ 1), this long tail acquires an exponential cutoff with slope 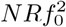, reflecting the new maximum size of a double-mutant lineage (*f*_*AB*_ ≲ 1*/NR*). Finally, when 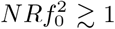, the distribution approaches the quasi-linkage equilibrium limit, with a sharp peak around Λ ≈ 1. **(B)** The total probability of observing all four haplotypes (Λ > 0) in a sample of size *n* = 10^4^ when *f*_*A*_ = *f*_*B*_ = *f*_0_. Symbols denote the results of forward-time simulations for neutral mutations with *N* = 10^6^ and *R* = 10^−1^, while the solid line denotes the theoretical prediction from Eq. (20).

In the opposite case where 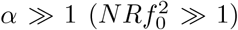, the distribution of Λ becomes sharply peaked around one, with a variance of order 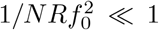. In this quasilinkage equilibrium (QLE) regime, recombination is so frequent that many double-mutant lineages are always present in the population at the same time. The sum of their individual sizes gives rise to the Gaussian-like behavior in Eq. (19) (Fig. 6A). This illustrates how higher moments of Λ can provide information about the transition to QLE, even when the average value of 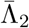 remains constant.

We can quantify the transition between these two regimes by examining the total probability of observing all four haplotypes (Λ > 0) as a function of the allele frequency scale *f*_0_. In Appendix G, we show that for a sample of size *n*, this probability can be approximated by

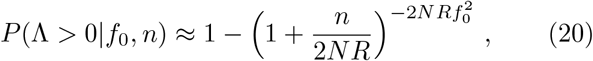

which grows quadratically with 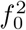 at low frequencies and approaches one when 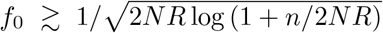 (Fig. 6B). The location of this transition provides another way to estimate the magnitude of *NR*.

When recombination is rare (*NRf*_0_ ≲ 1), the local equilibrium in Eq. (18) breaks down, which makes it difficult to obtain an analagous expression for the conditional distribution of Λ. Nevertheless, our heuristic calculations above suggest that rare double mutants that reach frequencies of order *f*_0_ will create a long tail in the Λ distribution, with Λ values as large as ∼ 1*/f*_0_ (see Fig. 6A). This long tail is balanced by an even smaller probability of reaching this maximum size 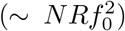, which brings the average back down to *NRf*_0_, consistent with our previous results in Eq. (10).

### Relaxing the assumption that both alleles are rare

All of the above results were derived for the first class of weighting functions in Eq. (3), which conditions on scenarios where both alleles are rare (*f*_*A*_, *f*_*B*_ ≲ *f*_0_ ≪ 1). We now consider extensions to the “single-rare” case in Eq. (5), where one of the two alleles can reside at a much larger frequency (*f*_*A*_ ≲ *f*_0_ ≪ *f*_*B*_).

Our solution for the “double-rare” case relied on a branching approximation for the three mutant haplotypes, which breaks down if one of the alleles (e.g. *B*) drifts to intermediate frequencies. However, if the *A* allele is present at a much lower frequency than *B*, then the frequency of the *B* allele will remain approximately constant over the lifetime of the *A* allele (Fig. 7). This separation of timescales suggests that we can analyze a simpler model where only the frequencies of *Ab* and *AB* haplotypes are changing. Interestingly, this two-locus problem is equivalent to a single-locus model of a subdivided population, where the demes correspond to the *B/b* alleles, and migration occurs when an *A* allele recombines onto a different *B/b* background. This yields a second branching approximation for the *Ab* and *AB* haplotypes, which allows us to obtain a solution for the conditional generating function 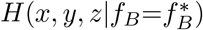 in Eq. (9) when *f*_*A*_ ≪ *f*_*B*_ (Appendix F).

**FIG. 7.**
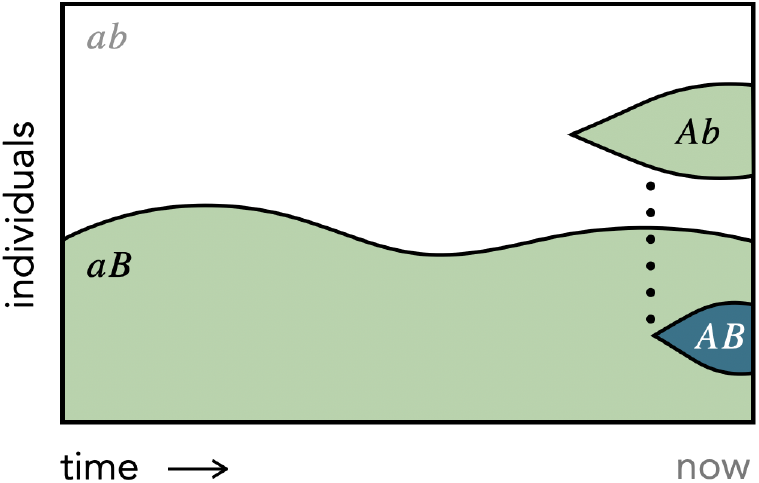
Schematic of the lineage dynamics contributing to homoplasy when one allele frequency is much larger than the other. The frequency of the common allele (*B*) remains effectively constant, while the dynamics of the rare allele (*A*) are much faster. In this regime, recombination events can be modeled as migrations between two approximately constant-sized demes (Appendxi F).

By applying these results to the homoplasy statistic in Eq. (6), we find that the average value 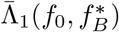 is qualitatively similar to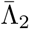 (Fig. 8). In the absence of selection or recurrent mutation, 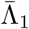 is again determined by the compound parameter *NRf*_0_,

**FIG. 8.**
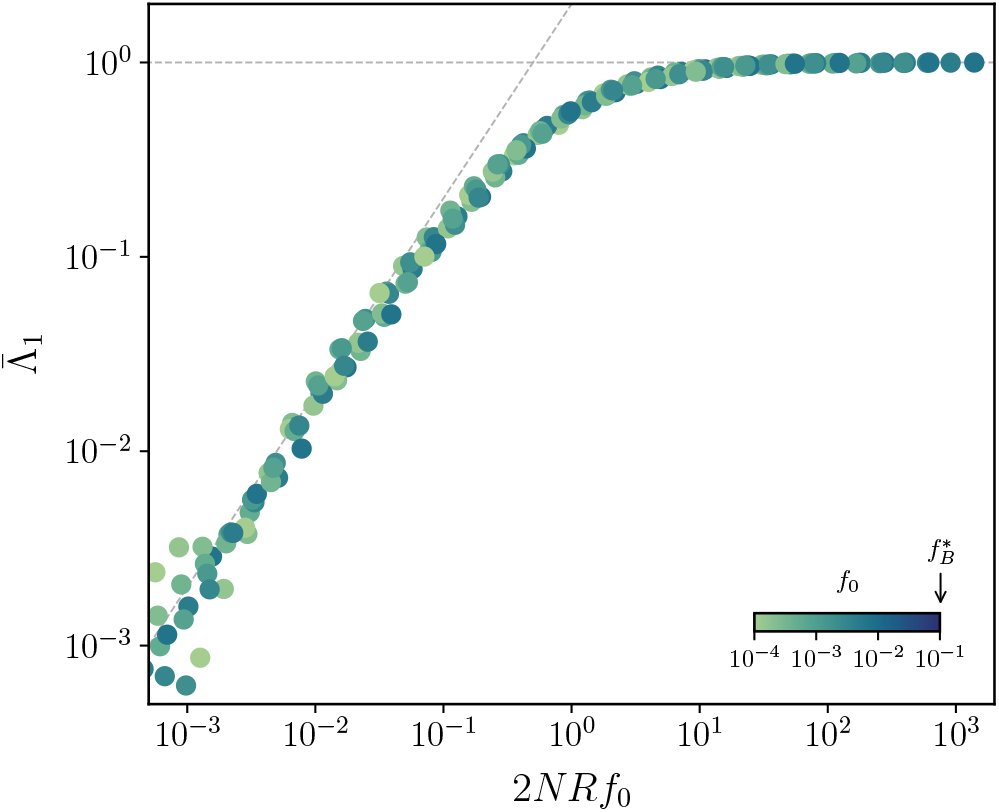
Frequency-resolved homoplasy,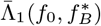 when only one of the alleles is rare. An analogous version of Fig. 2 illustrating the “single-rare” statistic in Eq. (6) with 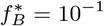 and *d* ∼ 10^−3^. Symbols denote the results of forward-time simulations for the same parameters as in Fig. 2, while the dashed lines denote the asymptotic predictions in Eq. (21).

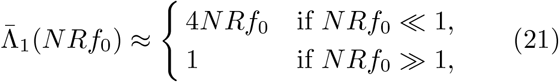

which transitions from a linear regime at low recombination rates (*NRf*_0_ ≪ 1) to a saturated regime when *NRf*_0_ ≫ 1, and is independent of 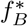.

When recombination is frequent (*NRf*_0_ ≫ 1), we can again derive an analytical expression for the distribution of Λ (Appendix F). In this “single-rare” case, the conditional distribution of Λ can be expressed as

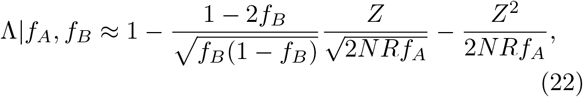

where *Z* is a Gaussian random variable with mean zero and variance one. When *NRf*_*A*_ ≫ max {1, 1*/f*_*B*_}, this distribution is sharply peaked around Λ ≈1. This implies that when the *B* allele is common (*f*_*B*_ ≳ 10%), the transition to the QLE regime occurs when *NRf*_0_ ≳ 1. The location of this transition is dramatically different from the case where both alleles were rare (Fig. 6), which required the stronger condition that 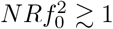.

Our heuristic picture provides an intuitive explanation for this difference. When *B* allele is present at intermediate frequencies, the rate at which the doublemutant lineages are created via recombination is of order *NRf*_*A*_*f*_*B*_ ∼*NRf*_0_, rather than 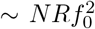. This implies that multiple *AB* recombinants will start to be produced when *NRf*_0_ ≫ 1, making it easier to attain linkage equilibrium.

## DISCUSSION

Homoplasy is a fundamental signature of recombination, but its quantitative behavior is less well understood. Here, we have studied a particular class of two-locus homoplasy statistics in a Wright-Fisher model under the joint action of recombination, recurrent mutation, additive and epistatic fitness costs, and genetic drift. By modeling the forward-time dynamics of the underlying lineages, we derived analytical expressions that predict how these homoplasy measures scale with the evolutionary parameters, as well as the present-day frequencies of the two alleles. We observed striking transitions as a function of this frequency scale, providing an independent lever for probing the dynamics of recombination across a range of ancestral timescales.

The homoplasy measures we considered in this work are conceptually similar to the *γ* statistic previously analyzed by Hey and Wakeley (1997), which represents the conditional probability of observing all four haplotypes in a sample of size *n* = 4. However, our focus on rare alleles allowed us to extend this approach to arbitrarily large sample sizes, and also to study the effects of negative selection and recurrent mutation that are challenging to model with traditional coalescent approaches (Wakeley, 2008; Walczak *et al*., 2012). The ability to condition on allele frequencies turned out to be particularly useful in these cases, suggesting new ways to potentially distinguish between these confounding evolutionary forces (Fig. 5).

Our homoplasy measures also share some qualitative features with traditional LD metrics like *r*^2^. For example, the complement of our homoplasy statistic, 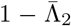, decays as ∼ 1*/NR* (Appendix E), similar to the frequency-weighted average of *r*^2^ (Good, 2022). But despite these high-level similarities, there are also important quantitative differences between these statistics that can be useful for distinguishing the underlying evolutionary forces. For example, previous work has shown that the numerical values of *r*^2^ at low recombination rates are strongly influenced by negative selection and genetic drift (Good, 2022), making it difficult to calibrate the overall scale. If there is greater epistasis among physically co-located sites (e.g. within the same gene or protein domain), then it is even possible to observe a decaying *r*^2^ curve – a classic signature of recombination – in an otherwise purely clonal population. In contrast, our analysis above shows that the values of 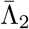 that are most informative about the underlying recombination rate 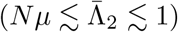 cannot be produced by other forces. Moreover, the transition to this regime occurs not only for loci with small map distances (i.e. *R* ≲ 1*/N*), but also for alleles with low present-day frequencies (*f*_0_ ≲ 1*/NR*). Since these rare alleles typically comprise the majority of segregating variants, these frequency-resolved homoplasy measures could be particularly useful for exploring the dynamics of genetic linkage in large genomic datasets.

As an illustrative example, we used this approach to measure frequency-resolved homoplasy in a collection of *n* = 4, 872 metagenomically-assembled genomes of the commensal human gut bacterium *Eubacterium rectale* (Fig. 9; Appendix H). Previous estimates of LD in this species have suggested that *E. rectale* strains in different hosts experience high rates of homologous recombination (Good, 2022; Liu and Good, 2024), which provides a natural opportunity for exploring the homoplasy metrics discussed above. The core genomes of these strains contained a total of 338,669 synonymous single-nucleotide variants (SNVs), the vast majority of which were rare (median minor allele frequency of 0.6%). We developed an unbiased estimator of the 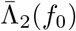 statistic in Eq. (4) that accounts for finite sample effects and applies for frequency scales as small as *f*_0_ ≳ 2*/n* (Appendix G). With a sample size > 4, 000, this dataset allowed us to quantify the emergence of homoplasy in *E. rectale* across nearly three orders of magnitude of allele frequencies (Fig. 9).

**FIG. 9.**
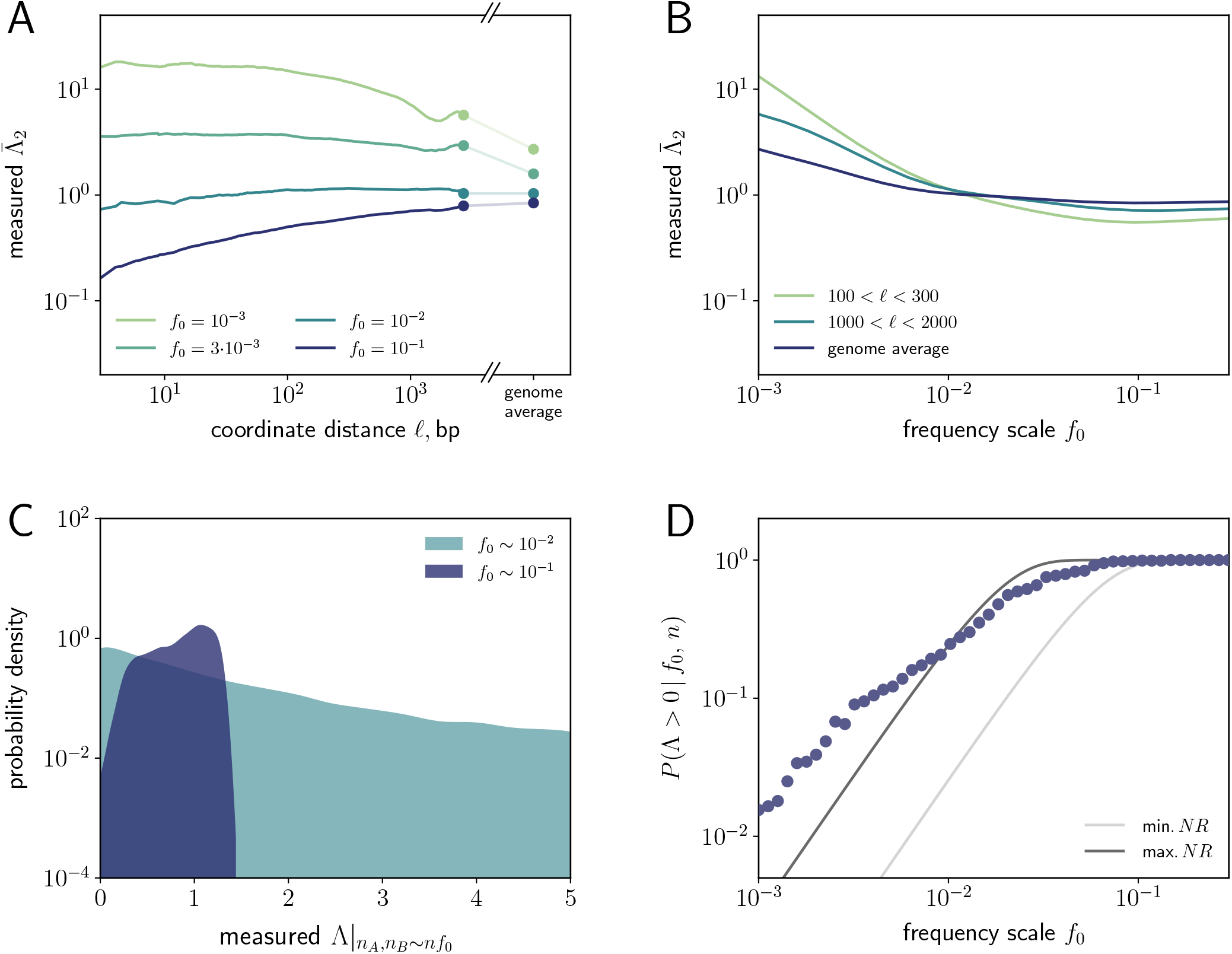
Frequency-resolved homoplasy in the commensal human gut bacterium *Eubacterium rectale*. SNVs were obtained for a sample of 4, 872 metagenomically assembled genomes reconstructed from different human hosts (Almeida *et al*., 2021; Appendix H). **(A)** Observed values of 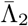 as a function of the estimated coordinate distance (*ℓ*) for pairs of synonymous SNVs in core genes. Solid lines were obtained by applying the unbiased estimator in Appendix G to all pairs of SNVs within 0.2 log units of *ℓ* in sliding windows. The genome-wide averages were calculated from randomly sampled pairs of SNVs from widely separated genes; the two estimates are connected by a faint line for visualization. **(B)** An analogous version of the top left panel as a function of the frequency scale *f*_0_. The observed values of 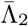grow larger as *f*_0_ decreases, contrary to the theoretical prediction from Fig. 2. **(C)** The observed distribution of Λ when both alleles are rare. The histograms show kernel density estimates for pairs of SNVs separated by *ℓ* > 10^6^ bp with marginal mutation frequencies *f*_0_ ∈ (0.175, 0.1925) (purple) and *f*_0_ ∈ (0.025, 0.0275) (teal). The shapes of these two distributions are qualitatively similar to the 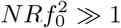 and 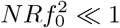 regimes predicted in Fig. 6A. **(D)** The total probability of observing all four haplotypes (Λ > 0) in a sample of size *n* = 4, 600 as a function of the frequency scale *f*_0_. Symbols denote the observed values computed for pairs of sites with allele frequencies *f*_*A*_, *f*_*B*_ in the range (0.3*f*_0_, 3*f*_0_), which were separated by *ℓ* > 10^6^ bp. The lines denote the theoretical predictions from Eq. (20) for the maximum and minimum possible values of *NR* inferred from the *f*_0_ values in panel C (30 ≲ *NR* ≲ 1, 500). At low frequencies, the observed value of *P* (Λ > 0|*f*_0_, *n*) is much larger than theoretically predicted.

At intermediate frequency scales (*f*_0_ ≳ 10^−2^), we found that the observed values of 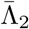 were qualitatively consistent with our theoretical predictions in Fig. 2: 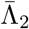 increases with the coordinate distance *ℓ* between the sites (a proxy for their total map distance *R*), and eventually saturates at one at large distances (Fig. 9A). Similarly, the conditional distribution of Λ for the most widely separated sites (*ℓ* ≳ 10^6^ bp) exhibits a transition from a broad distribution at lower frequencies (*f*_0_ ≈ 10^−2^) to a unimodal shape when *f*_0_ ≈ 10^−1^ (Fig. 9C). This shift is qualitatively consistent with the predicted transition to the quasi-linkage equilibrium regime 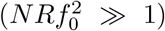 in Fig. 6A. Since the typical SNV in *E. rectale* has a frequency < 1%, these results indicate that the vast majority of SNVs have not yet reached linkage equilibrium, even though the genome-wide values of LD are low (Garud *et al*., 2019). This observation has important implications for the application of demographic inference methods like *∂*a*∂*i (Gutenkunst *et al*., 2009; Kim *et al*., 2017; Mah *et al*., 2023), which assume that most variants are in linkage equilibrium with each other.

In addition to these qualitative similarities, we also observed several striking departures from the predictions of our simple model above. For example, at sufficiently low frequencies (*f*_0_ ≲ 10^−2^), we find that 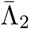 decreases as a function of *f*_0_ (Fig. 9B), in contrast to what we would expect from Fig. 2. The reason for this discrepancy can be traced to the distribution of Λ in Fig. 9C: while the nonzero part of this distribution is qualitatively similar to our predictions in Fig. 6A, the total probability of observing a nonzero value increases more slowly with *f*_0_ than expected theoretically (Fig. 9D). This implies that there are more combinations of all four haplotypes at lower frequencies than we would expect in our model, which is responsible for elevating the mean value 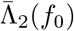 in Fig. 9B. This illustrates how a quantitative understanding of homoplasy can reveal qualitative features of the data that require additional theoretical explanation. The existence of such discrepancies is not too surprising, since we have focused on a simple evolutionary model that omits many known complexities of natural microbial populations. One important limitation is the assumption of a panmictic population with a constant size. In reality, host-associated organisms like gut bacteria can exhibit complex population structures that depend on their history of dispersal and co-diversification with their hosts (Falush *et al*., 2003; Mah *et al*., 2023; Suzuki *et al*., 2022). Another crucial assumption is the absence of positive selection and hitchhiking of linked neutral loci, which are thought to play an important role in shaping the genetic diversity of natural bacterial populations (Birzu *et al*., 2023; Liu and Good, 2024; Wolff and Garud, 2023). While further work will be required to account for these effects, many of the qualitative features of our analysis – in particular, the lineage decomposition in Fig. 1 – will continue to apply in these more complex scenarios. Our theoretical framework may therefore provide a useful starting point for understanding frequency-resolved linkage more broadly.

## DATA AVAILABILITY

Source code for forward-time simulations, numerical calculations, data analysis, and figure generation is available on Github (https://github.com/alyulina/linkage-equilibrium). Polymorphism data from *E. rectale* were obtained from a previous study (Almeida *et al*., 2021) and can be accessed using the accessions listed in that work.

## ACKNOWLEDGEMENTS

We thank Noah Rosenberg, Dmitri Petrov, Jeffrey Spence, and members of the Good lab for fruitful discussions and feedback on the manuscript. This work was supported in part by the Alfred P. Sloan Foundation (FG-2021-15708), NIH NIGMS (R35GM146949), and a Stanford Bio-X Bowes Fellowship (to Z.L.). B.H.G. is a Chan Zuckerberg Biohub – San Francisco Investigator.

## Appendix A Forward-time simulations

We validated our analytical predictions by comparing them to forward-time simulations of the two-locus Wright-Fisher model. In each generation *t*, the haplotype frequencies were first updated using the deterministic update rule

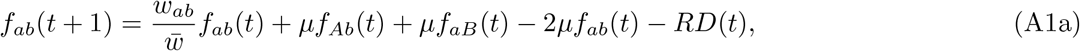

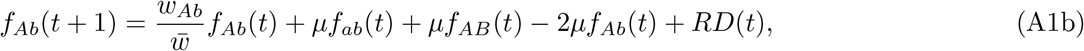

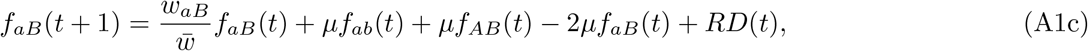

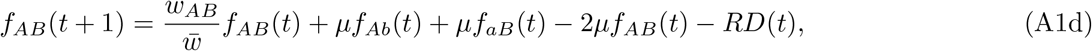

where *D* ≡ *f*_*AB*_*f*_*ab*_ − *f*_*Ab*_*f*_*aB*_ is the coefficient of linkage disequilibrium, *w*_*i*_ ≡ exp(*s*_*i*_) is the Wrightian fitness of the haplotypes, and 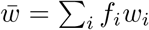 is the mean fitness of the population. Genetic drift was then incorporated by drawing a random number of individuals for each haplotype using a Poisson distribution with mean *Nf*_*i*_.

To speed up the simulations, we only simulated time intervals where both loci contained a segregating mutation. Without loss of generality, we let *A* correspond to the earlier of the two mutations, and *B* correspond to the later one. We assumed that when the *B* mutation arises, the initial frequency of the *A* mutation can be approximated by the single-locus site frequency spectrum (Sawyer and Hartl, 1992),

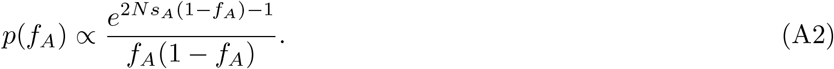

This will be a good approximation as long as *Nμ* ≪ 1. Based on this random value of *f*_*A*_, the new *B* mutation was assigned to an *AB* or *aB* haplotype with probabilities *f*_*A*_ and 1 − *f*_*A*_, respectively. The resulting population was then evolved using the update rule above until one of the two mutations went extinct, and the process was then restarted with a new pair of mutations. The frequencies of the four haplotypes were recorded every Δ*t* = 100 generations, and were used to generate the figures in the main text.

## Appendix B Perturbative solution for the moment generating function of the haplotype frequency distribution

Good (2022) previously derived a perturbative solution for the moment generating function in Eq. (7) that is valid for sufficiently small allele frequencies. We reproduce this solution here for completeness, since it will form the basis for many of our analytical calculations below.

Since the generating function does not explicitly depend on the allele frequencies, it is helpful to define a rescaled version of Eq. (7),

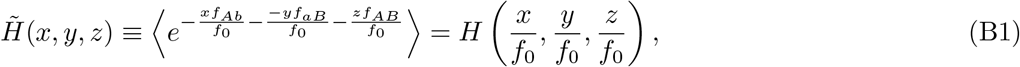

which is dominated by frequencies ≲ *f*_0_ when *x, y*, and *z* are 𝒪(1). Choosing small values of *f*_0_ then allows us to focus on small values of *f*_*A*_ and *f*_*B*_.

When *f*_*A*_ and *f*_*B*_ are both small compared to one (*f*_0_ ≪ 1), the two-locus model in Appendix A reduces to the branching-process-like form,

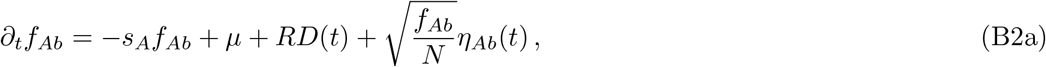

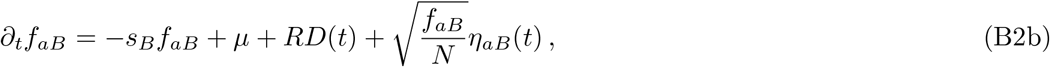

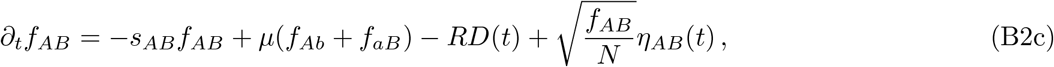

where *D* ≡ *f*_*AB*_ − *f*_*A*_*f*_*B*_ and *η*_*Ab*_(*t*), *η*_*aB*_(*t*), *η*_*AB*_(*t*) are independent Brownian noise terms with mean zero and variance one (Good, 2022). By differentiating Eq. (B1) with respect to time and applying the stochastic dynamics in Eq. (B2), one finds that the generating function must satisfy the partial differential equation,

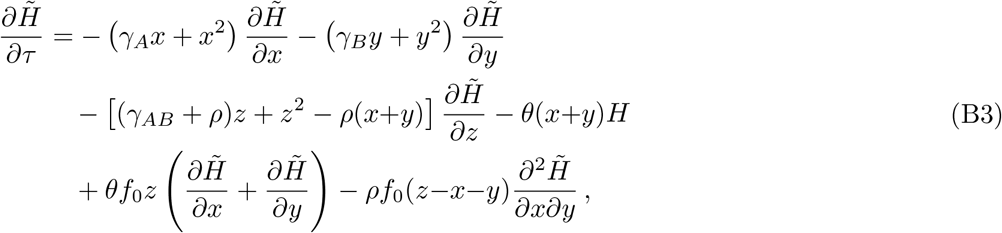

subject to the initial condition 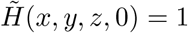, where we have defined a collection of scaled variables,

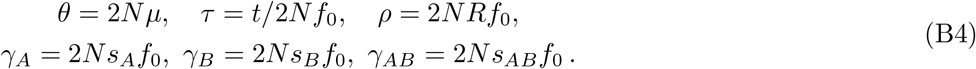

In the limit that *θ* ≪ 1 and *f*_0_ ≪ min {1, *ρ*^−1^}, the solution to Eq. (B3) can be expressed as a power series (Good, 2022):

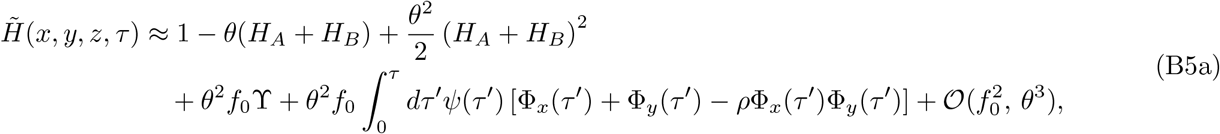

where

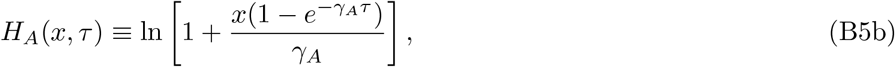

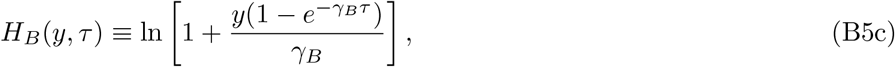

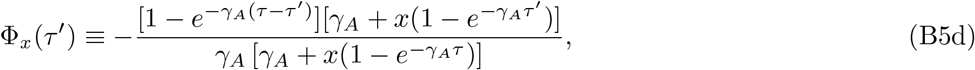

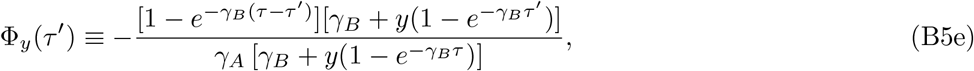

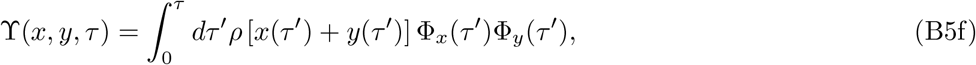

and *ψ*(*τ* ′) is a solution to the characteristic curve,

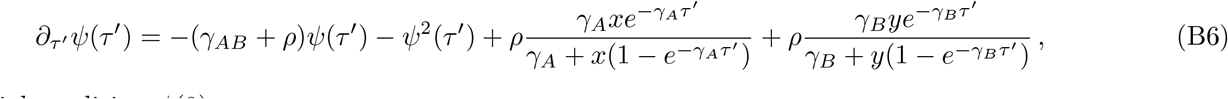

with the initial condition *ψ*(0) = *z*.

The individual terms in the perturbation expansion in Eq. (B5a) correspond to different classes of haplotype frequency trajectories. For example, the *H*_*A*_ and *H*_*B*_ terms enter the series at first order in the scaled mutation rate (*θ* = 2*Nμ*) and correspond to trajectories where only one of the two loci is mutated at any given time. Two-locus statistics like Λ that require both sites to be mutated will therefore only start to enter at order 𝒪(*θ*^2^). The functional form of the 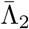 statistic in Eq. (4) leads to further simplifications. The numerator of Eq. (4) depends on a triple derivative of the generating function,

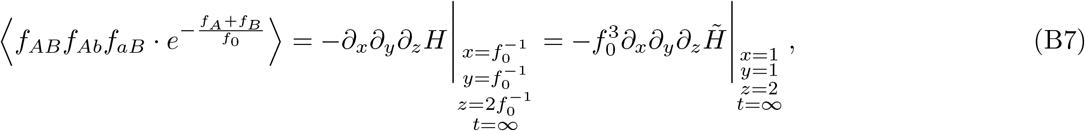

which we can evaluate using Eq. (B5). Many of the terms in Eq. (B5a) are independent of z, and therefore vanish when taking the *z* derivative in Eq. (B7). The lowest-order terms that depend on *z* are the *ψ*(*τ* ′) terms, so that Eq. (B7) reduces to

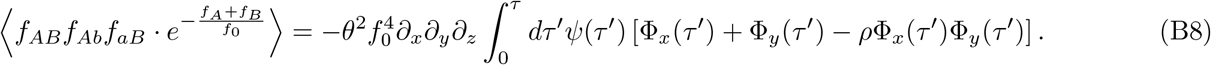

In this way, the problem of calculating 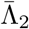 reduces to finding the solution of the characteristic curve in Eq. (B6). Solutions to this nonlinear equation were previously obtained by Good (2022) for the special case where *x* = *y* = 1. However, the presence of the *x* and *y* derivatives in Eq. (B8) now require us to extend this solution to arbitrary values of *x* and *y* in the local neighborhood of *x* = *y* = 1, where the previous solution method employed by Good (2022) breaks down.

Here we account for this behavior using two complementary approaches. In Appendix C, we first outline a numerical method for directly calculating the derivatives of *ψ*(*τ* ′) with respect to *x, y*, and *z*. We use this approach to calculate the numerical curves in Figs. 2, 3, & 5. In addition, we also use a separation of timescales approximation to derive approximate analytical solutions to Eq. (B6) that apply for specific parameter regimes, which correspond to cases where selection and recombination are weak compared to drift (Appendix D), or strong compared to drift (Appendix E), respectively. These asymptotic solutions cover a broad range of parameter space, and provide additional insights into the evolutionary dynamics in each regime.

## Appendix C Numerical solution for 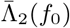

The characteristic curve in Eq. (B6) is difficult to solve in the general case because the inhomogeneous terms vary over many different timescales. However, we saw in Eq. (B8) that the moments of Λ only depend on the behavior of *ψ*(*τ* ′) in the local neighborhood around *x* = 1, *y* = 1, and *z* = 2, suggesting a perturbative expansion of the form

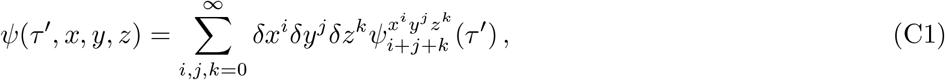

where *δx* = *x* − 1, *δy* = *y* − 1, and *δz* = *z* − 2.

After substituting the ansatz in Eq. (C1) into Eq. (B6), expanding in powers of *δx, δy*, and *δz*, and collecting like terms, we obtain a system of ordinary differential equations for the 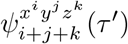 functions,

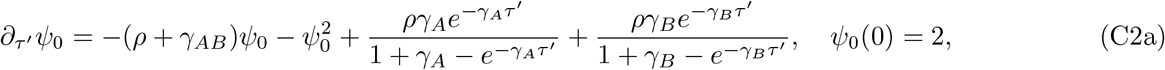

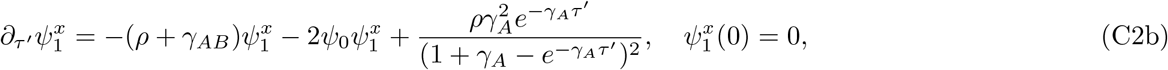

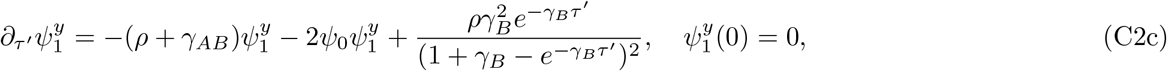

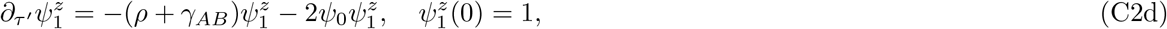

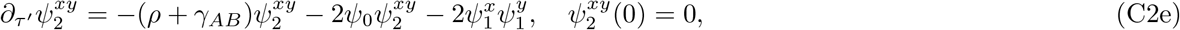

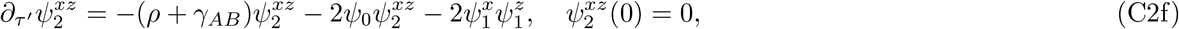

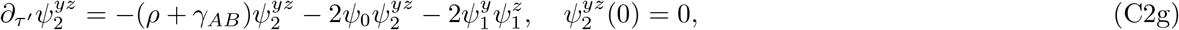

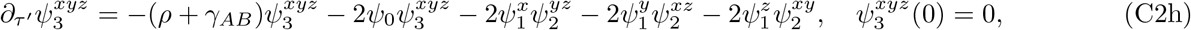

which are independent of *δx, δy*, and *δz*. We solved this system numerically using the solve ivp() function from the SciPy Python library (Virtanen *et al*., 2020).

We combined these numerical solutions with Eq. (B8) to compute our frequency-resolved homoplasy statistic 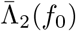. Substituting Eq. (C1) into Eq. (B8), we find that the leading order contribution to the numerator of 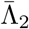 is given by

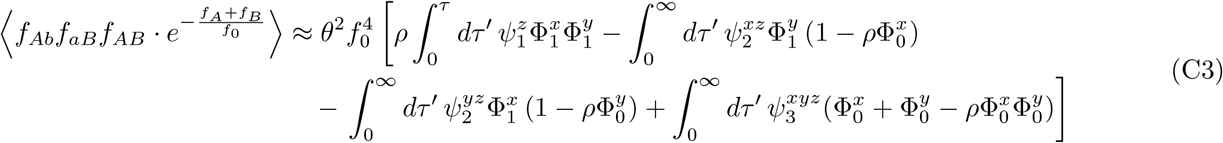

where we have defined

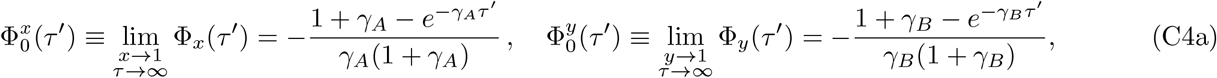

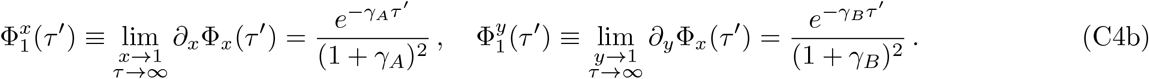

At lowest order in *f*_0_, the denominator of 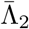 will usually be dominated by the two single mutation terms, so that

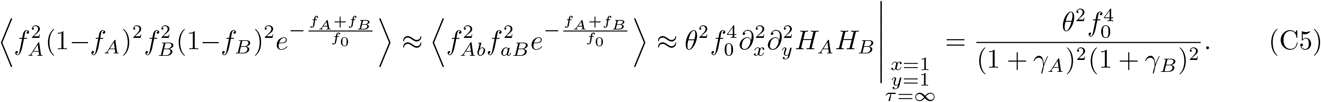

Combining this with Eq. (C3), we can obtain a corresponding expression for 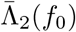:

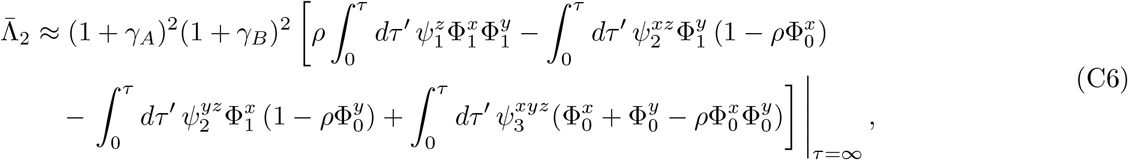

as a function of the numerical solutions 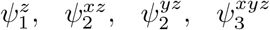 from Eq. (C2). In the neutral limit (*γ*_*A*_ → 0, *γ*_*B*_ → 0, *γ*_*AB*_ → 0), this expression further reduces to

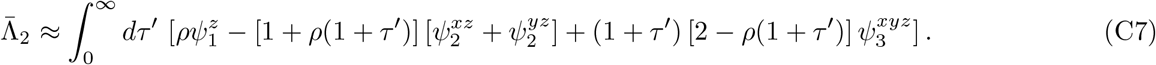

which only depends on the value of the compound parameter *ρ* = 2*NRf*_0_. We evaluated this integral numerically by approximating it as a Riemann sum over the discretized solutions for 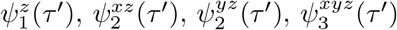 above using a step size of *δτ* ′ = 3 10^−6^*/ρ*. We used this procedure to generate the theoretical curves in Figs. 2, 3A, and 5A in the main text.

## Appendix D Analytical solution for 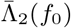 for neutral loci and weak recombination

To obtain an analytical solution for 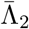, we begin by considering the limit where *γ*_*A*_, *γ*_*B*_, *γ*_*AB*_ are small compared to both 1 and *ρ*. Physically, this means that selection is weak in comparison to drift and recombination. Recall that since *γ*_*A*_, *γ*_*B*_, *γ*_*AB*_ contain a power of *f*_0_, this regime also applies to nominally deleterious alleles, provided that *f*_0_ is sufficiently small. In this case, we can rewrite Eq. (B6) as

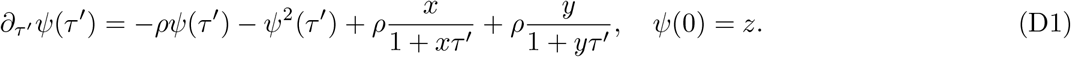

If we further assume that *ρ* ≪ 1, we can solve Eq. (D1) perturbatively in powers of *ρ*, treating the recombination terms as a correction to the otherwise asexual dynamics. In the absence of recombination, the characteristic curve in Eq. (D1) reduces to a logistic equation, whose solution is given by

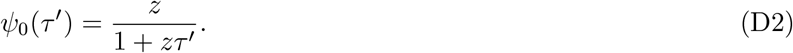

Corrections to this zeroth-order solution can be found by considering the series ansatz

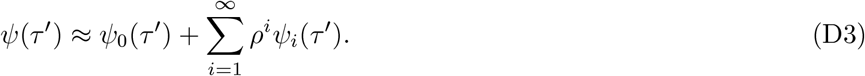

Substituting the above series expansion into Eq. (B6) and matching the coefficients in front of powers of *ρ*, we obtain for the first-order correction

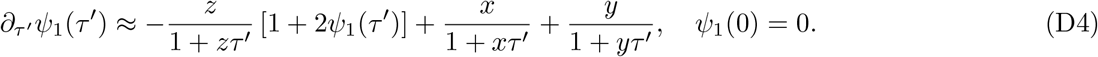

We can solve this equation with the method of variation of constants, which yields

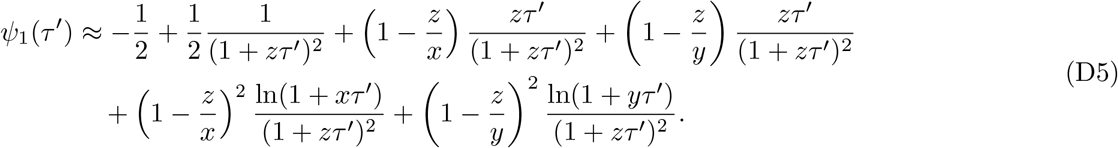

We are now in a position to find the averages in Eq. (4). To the lowest order in *ρ*, the numerator of 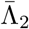 follows from Eq. (8) as

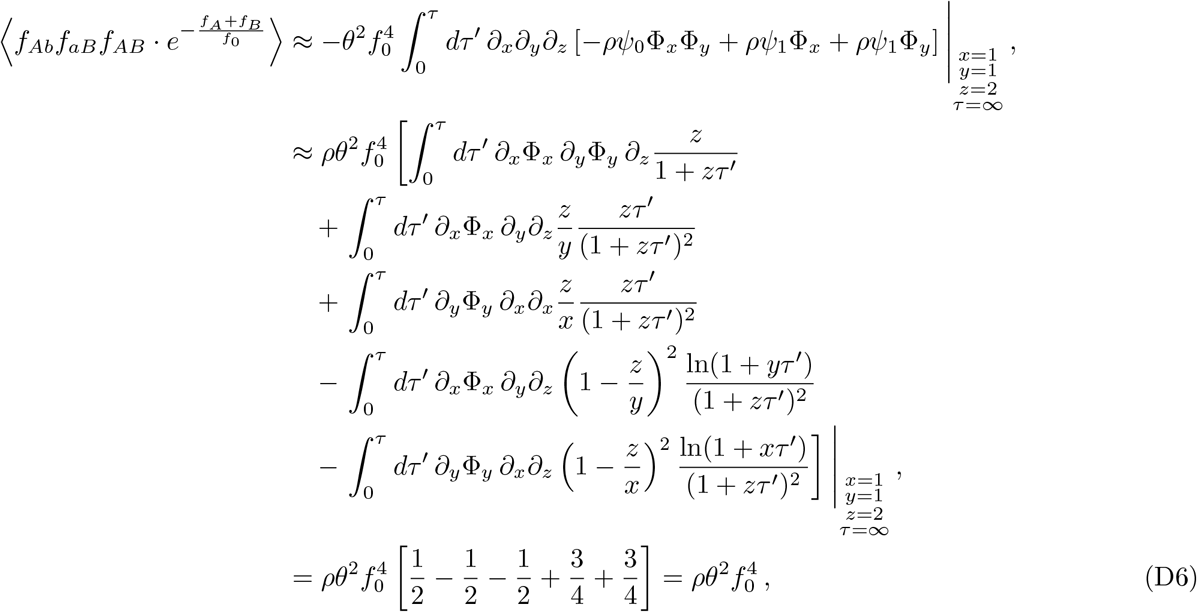

where have used the identities

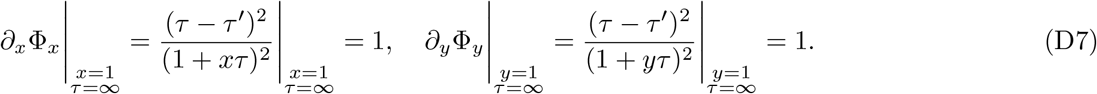

We note that terms in the integrand of Eq. (D6) correspond to the contributions coming from recombining separate (first term) and nested (the last two terms) mutations. This observation allows us to connect these terms with their diagrammatic representation in Fig. 1 and quickly estimate their contributions with a heuristic approach.

The dominant contribution to the denominator of 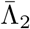 follows from Eq. (8) as

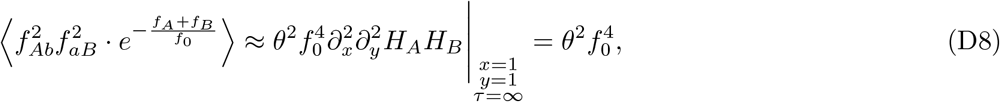

which derives from the term in Eq. (B5a) corresponding to two separate separate single mutants. Combining these two results together, we find that 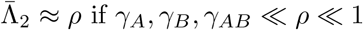.

## Appendix E Analytical solution for 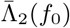 for strong selection or recombination

When selection or recombination are strong compared to drift, we can solve Eq. (B6) with the separation of timescales approach (Good, 2022), treating the drift term as a perturbative correction. In the limit that either *γ*_*AB*_ or *ρ* are large compared to one, we can rescale time in Eq. (B6) so that

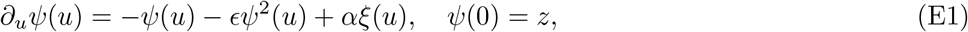

where *u* = *τ* ′*/ϵ* is the scaled time, *ϵ* = 1/(*γ*_*AB*_ + *ρ*), *α* = *ϵρ, β*_*A*_ = *ϵγ*_*A*_, *β*_*B*_ = *ϵγ*_*B*_, and

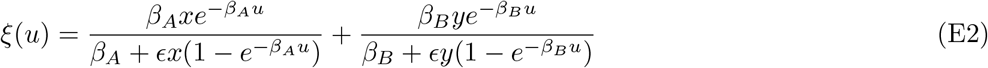

is a function independent of *z*.

We can solve Eq. (E1) using a perturbation expansion in *ϵ*, defining

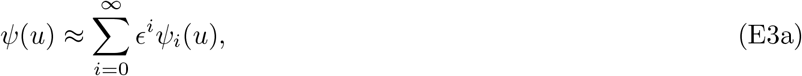

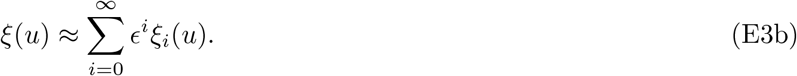

Substituting these expressions into Eq. (E1), we find that the zeroth order terms in *ϵ* satisify

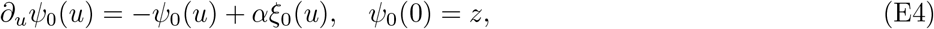

and hence

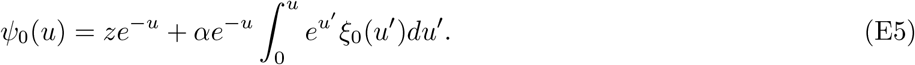

Likewise, the first-order contribution in *ϵ* satisfies

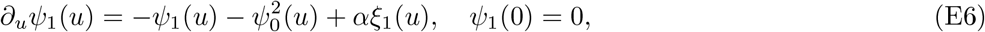

and hence

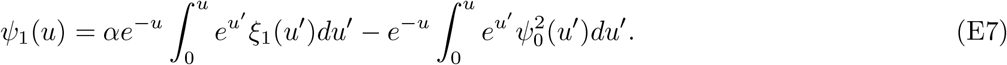

Finally, at the second order in *ϵ*, we have

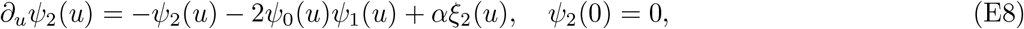

and hence

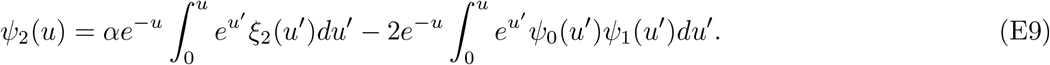

The *ξ*_*i*_ terms above will depend on the specific form of selection. We consider three different regimes below.

### Case 1: Both single mutants are strongly deleterious

In the case that *γ*_*A*_, *γ*_*B*_ ≫1, we can expand the *ξ*(*u*) function as

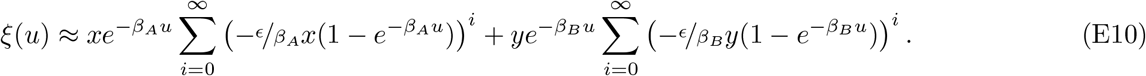

Substituting this expression into Eqs. (E5), (E7), and (E9) above and then applying Eq. (8), we find that the numerator of 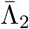 is given by

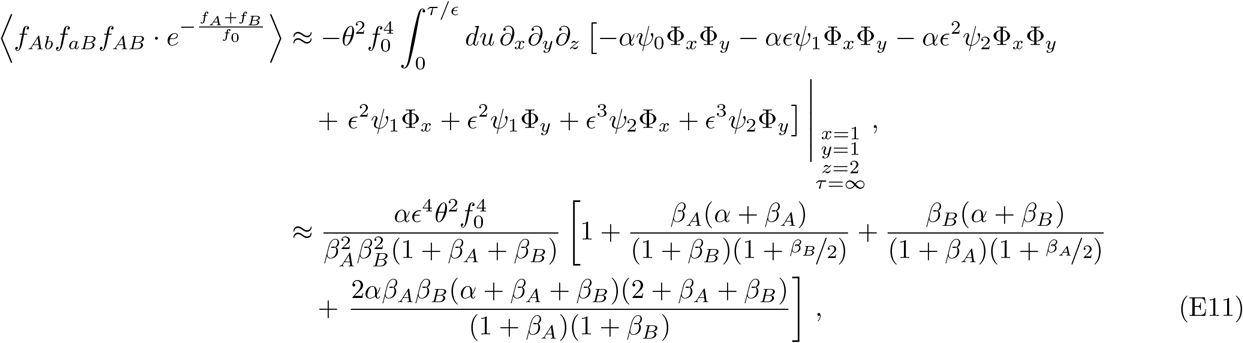

where we have used used

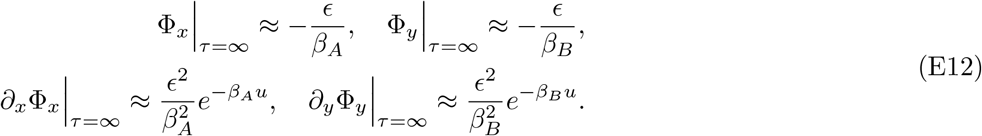

This result holds for any value of *ρ* if *γ*_*A*_, *γ*_*B*_ ≲ *γ*_*AB*_, and for small values of *ρ* ≫ 1 when *γ*_*A*_, *γ*_*B*_ ≫ *γ*_*AB*_.

As long as *γ*_*A*_, *γ*_*B*_ ≲ *γ*_*AB*_, the two separate single mutants will provide the dominant contribution to the denominator of 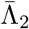. Therefore, the denominator follows from Eq. (8) as

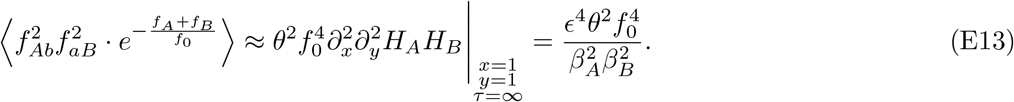

Dividing Eq. (E11) by Eq. (E13), we obtain

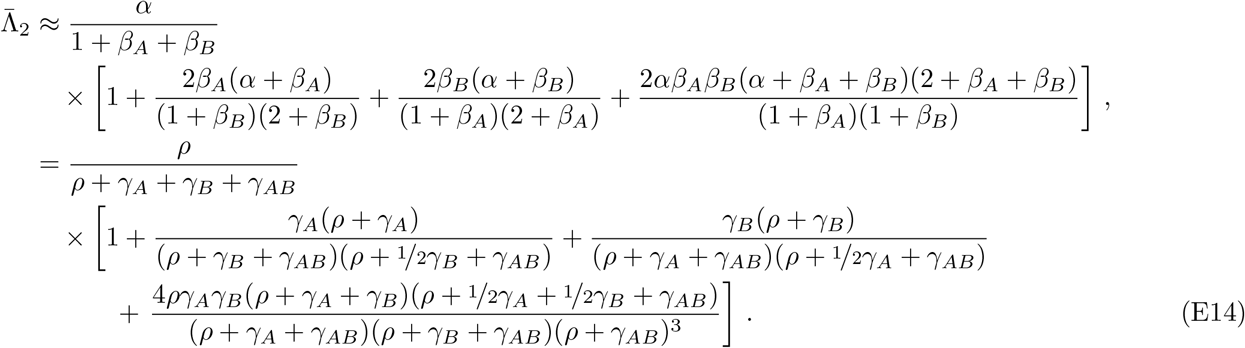

Eq. (E14) was used to generate the theory curves in Figs. 3, 4, & 5 in the main text. In the case of additive fitness effects (*γ*_*A*_ = *γ*_*B*_ = *γ* and *γ*_*AB*_ = 2*γ*), the expression in Eq. (E14) reduces to

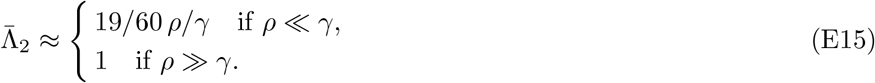

Defining the effective fitness cost as *s*_*e*_ = 30/19*s*_*AB*_ yields Eq. (14) in the main text.

### Case 2: Only one of the two mutations is strongly deleterious

In the limit that *γ*_*A*_ ≫ 1, *γ*_*B*_ = 0, we can solve Eq. (E1) by considering a different series expansion for *ξ*(*u*),

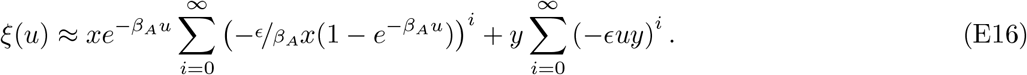

In this case, to the lowest order in *ϵ*, the numerator of 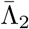 follows from Eq. (8) as

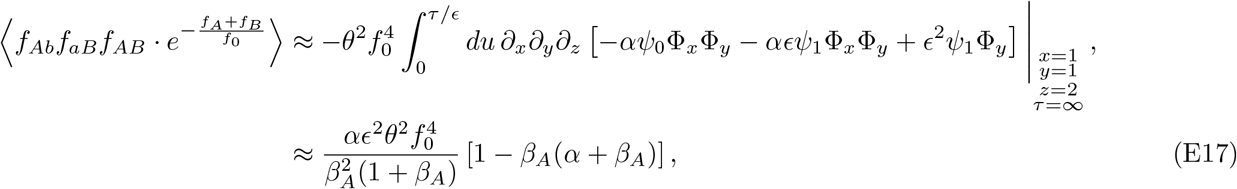

where we have used used

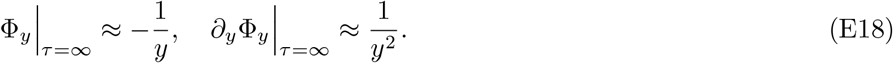

The denominator of 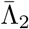 follows from Eq. (8) as

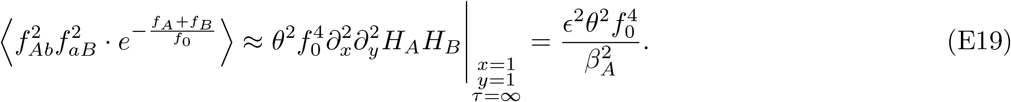

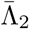 then follows as

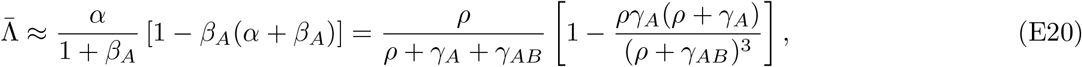

and for *γ*_*AB*_ = *γ*_*A*_ = *γ*,

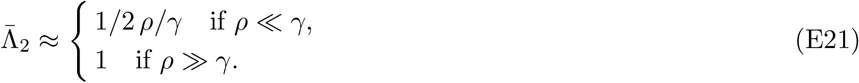

Defining the effective fitness cost as *s*_*e*_ = 2*s*_*AB*_ yields Eq. (14) in the main text.

### Case 3: Both single mutants are neutral

Finally, in the limit that both loci are neutral, but either recombination is strong or epistasis is strong, considering

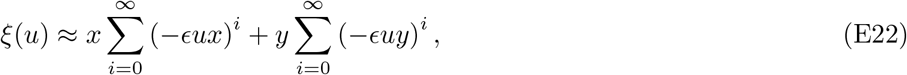

from Eq. (8) we obtain the numerator of 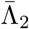 to the first order in *ϵ*,

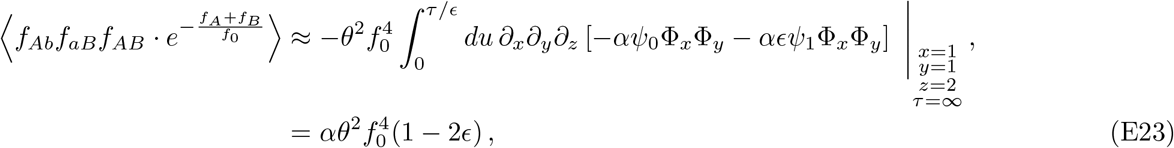

where we have used

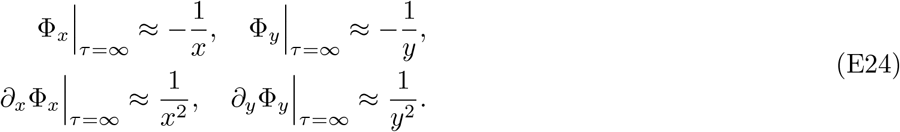

Approximating the denominator of 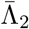 by Eq. (D8), we find that

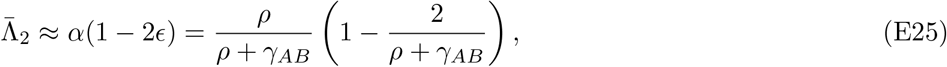

and therefore

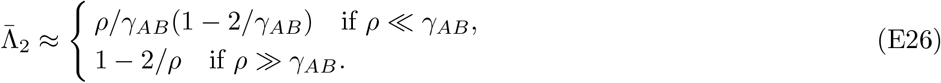

Defining the effective fitness cost as *s*_*e*_ = *s*_*AB*_ yields Eq. (14) in the main text.

## Appendix F Relaxing the assumption that both alleles are rare

When *f*_*A*_ ≪ *f*_*B*_, the two-locus dynamics in Appendix A can be approximated by a branching process model for the *A* mutation on timescales that are short compared to *Nf*_*B*_. Defining the rescaled frequencies *f*_1_ ≡ *f*_*Ab*_*/f*_*b*_ and *f*_2_ ≡ *f*_*AB*_*/f*_*B*_, these linearized dynamics can be written in the convenient form,

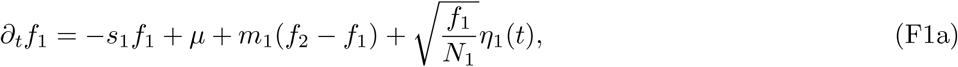

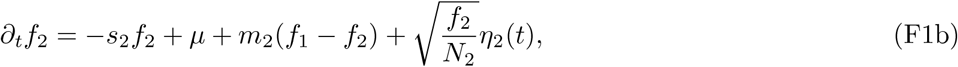

where we have defined a new set of constants *N*_1_ ≡ *Nf*_*b*_, *N*_2_ ≡ *Nf*_*B*_, and *m*_1_ ≡ *Rf*_*B*_, *m*_2_ ≡ *Rf*_*b*_. Writing the model in this way shows that the two-locus model maps on to a *single-locus* model with migration between two demes, in which both the migration rates *m*_1_, *m*_2_ and the effective population sizes *N*_1_, *N*_2_ depend on the frequency of the common *B* allele.

Rewriting Eq. (6) in terms of the rescaled variables yields an analogous relation for 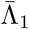,

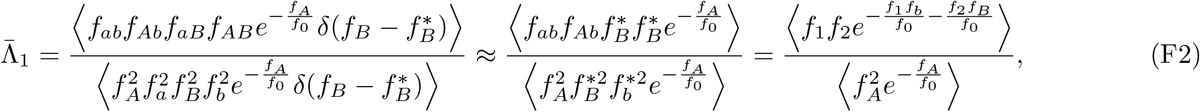

which can be calculated from the joint moment generating function for *f*_1_ and *f*_2_,

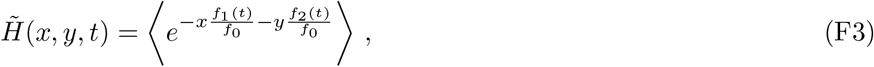

using the identity

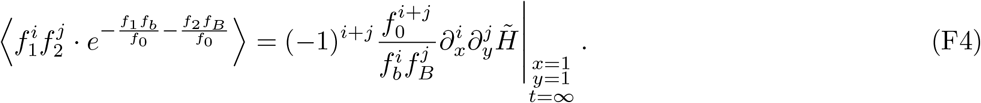

By differentiating Eq. (F3) with respect to time and applying the stochastic dynamics in Eq. (F1), we find that the generating function 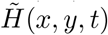 must satisfy the partial differential equation,

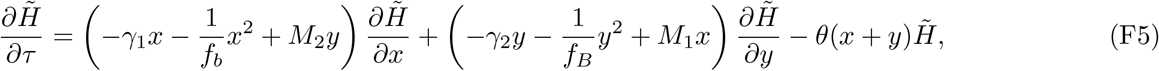

where we have defined the scaled parameters,

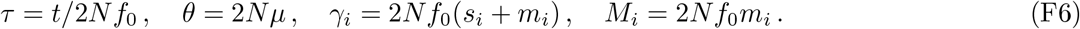

We derive approximate solutions to this equation in two different regimes below.

### Case 1: Rare migration/recombination

When *θ* ≪ 1 and *M*_*i*_ ≪ 1, we can obtain a perturbative solution for 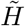 by repeating the perturbation calculation in Appendix B. To the lowest order in *M*_*i*_ and *θ*, we find that

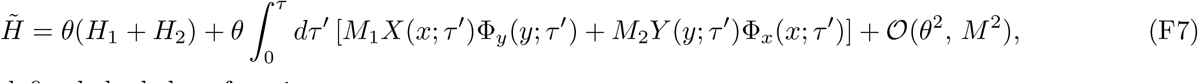

where we have defined the helper functions

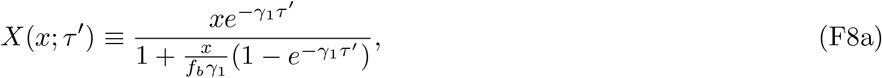

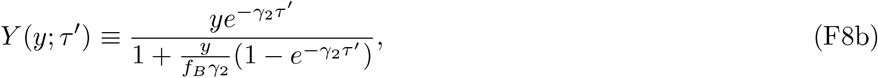

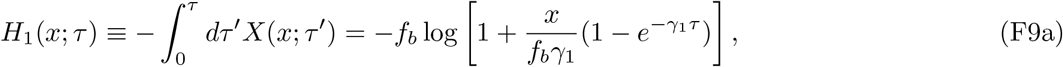

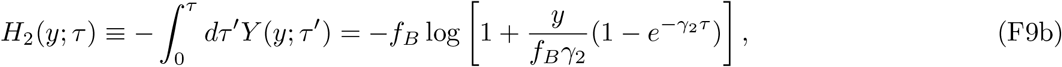

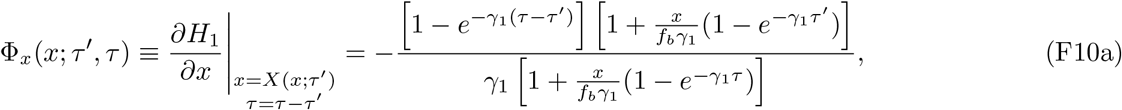

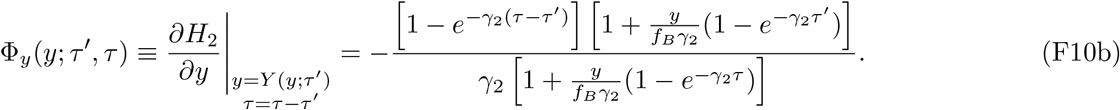

By combining this solution with Eqs. (F2) and (F4), we can obtain an analytical approximation for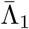.

Observing both *A* haplotypes in the population requires at least one *A* mutation event and one migration/recombination event. This means that only the 𝒪(*θ, M*_*i*_) term in Eq. (F7) contributes to the numerator of 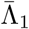:

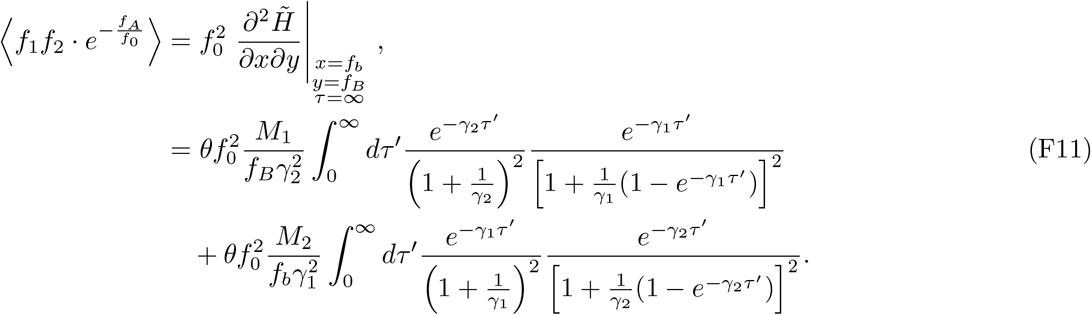

Since the integrals above involve two timescales, ∼ 1*/γ*_2_ and ∼ 1*/γ*_1_, it is difficult to find the solution analytically. However, since our perturbative expansion is only valid at the lowest order in *M*_*i*_, we can further expand the integrands in Eq. (F11) and evaluate them at the lowest order. For simplicity, we restrict our analysis to the case where both alleles are neutral. In this case, *γ*_1_ = *M*_1_ = *ρf*_*B*_, *γ*_2_ = *M*_2_ = *ρf*_*b*_. When *ρ* → 0, Eq. (F11) becomes

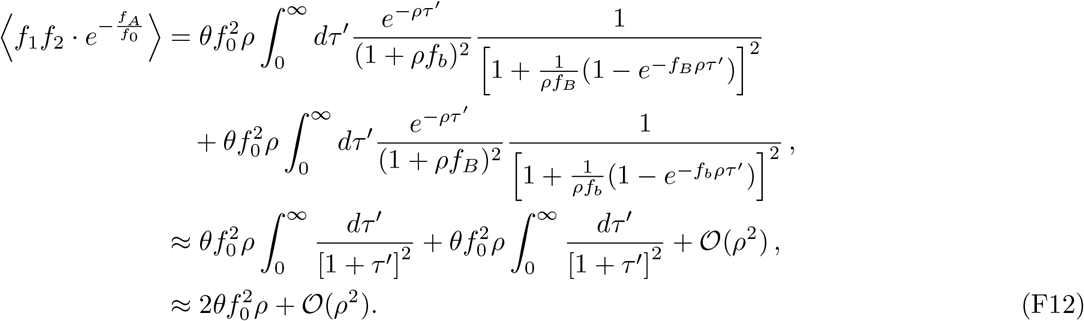

To calculate the denominator of 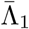, we simply recall that the neutral site frequency spectrum is given by

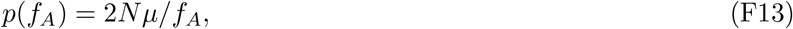

and therefore

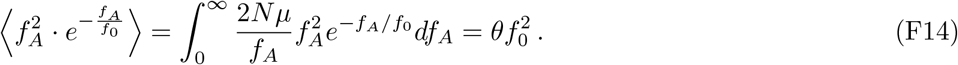

Combining the results above, we find that 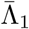 scales linearly with the rate of recombination,

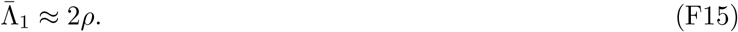

### Case 2: Frequent migration/recombination

In the regime where migration or recombination are frequent, we expect *f*_1_ and *f*_2_ to remain close to the average *f*_*A*_, which varies on the slower timescale ∼*Nf*_*A*_. This suggests that we can use a separation of timescales approach similar to Eq. (18) to model the fast dynamics of δ*f* ≡*f*_1_ − *f*_2_ conditioned on a fixed value of *f*_*A*_ ≡ *f*_1_*f*_*b*_ + *f*_2_*f*_*B*_. Subtracting the two equations in Eq. (F1) yields a corresponding equation for δ*f*,

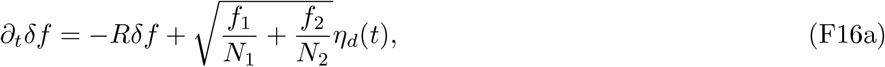

which reduces to

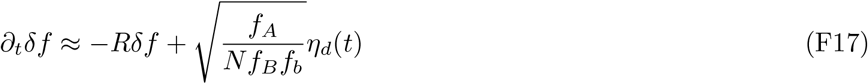

in the limit that δ*f* ≪ *f*_*A*_. For a fixed value of *f*_*A*_, these short-time dynamics attain the local equilibrium,

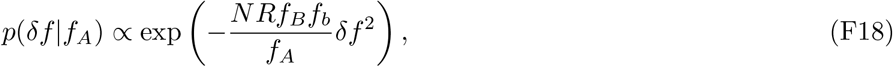

which is a Gaussian distribution with mean zero and variance 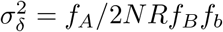.

We can use this result to obtain an analogous approximation for 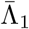. The numerator of Eq. (F2) is given by

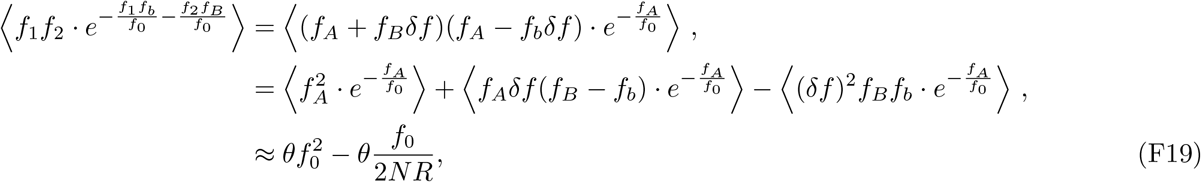

where in the last line we have first used Eq. (F18) to compute the averages over δ*f*, and then averaged over *f*_*A*_ using the neutral site frequency spectrum *p*(*f*_*A*_) = *θ/f*_*A*_. Combining this result with Eq. (F14) above, we find that

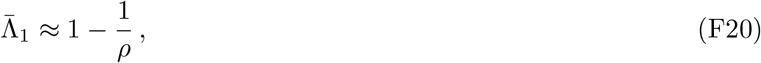

as expected.

We can also use Eq. (F18) to derive an approximation for the distribution of Λ in this regime. The relationship between Λ and δ*f* is given by

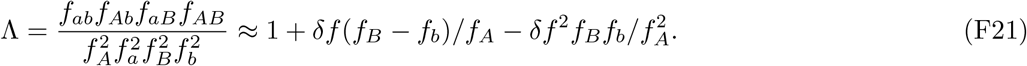

Since δ*f* follows a Gaussian distribution, we can rewrite the above expression as

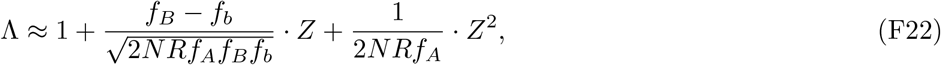

where *Z* is a Gaussian random variable with mean 0 and variance 1.

## Appendix G Estimating frequency-resolved homoplasy in finite samples

In order to connect our theoretical predictions for the moments of Λ with empirical observations, we need to account for the effects of finite sampling. In a sample of *n* genomes, we cannot directly observe the population haplotype frequencies (*f*_*ab*_, *f*_*Ab*_, *f*_*aB*_, *f*_*AB*_), but rather the discrete counts 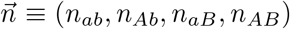. Although genomic datasets routinely exceed thousands of samples nowadays and will continue to expand in scale, sampling noise will remain important for rare alleles at low frequencies (*nf*_0_ ∼ 10).

### Finite-sample estimator for 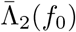

To accurately estimate averages such as 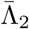 across a range of allele frequencies, we can rely on a class of unbiased estimators for frequency weighted moments that we have used in our earlier work (Good, 2022). This approach constructs a function,

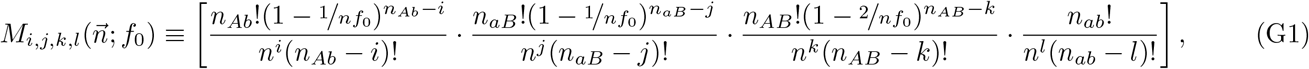

which has the property that when averaged over both the sampling noise and the noise from genetic drift, it is equal to the frequency-weighted average

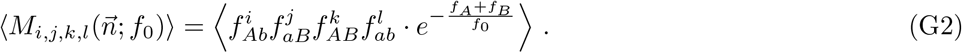

This means that we can estimate any frequency-weighted moment by aggregating many functionally similar pairs of genetic loci (e.g. similar recombination map length) and averaging the corresponding *M*_*i,j,k,l*_ across these pairs. For example, the numerator of 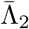 can be estimated as

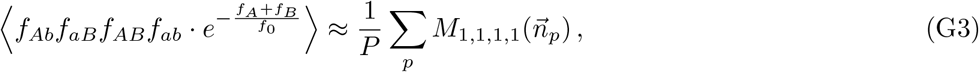

where *p* indexes a pair of loci and *P* is the total number of functionally similar pairs of loci.

The denominator of 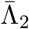,

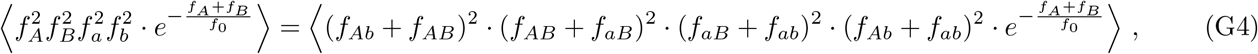

can be estimated in the same fashion using Eq. (G2). Since the the polynomial in Eq. (G4) involves a total of 2^8^ terms when expanded in powers of *f*_*ab*_, *f*_*Ab*_, *f*_*aB*_, *f*_*AB*_, we implemented a Python function that programmatically constructs the associated estimator for a given polynomial. Specifically, this function expands a polynomial using the Python package SymPy (Meurer *et al*., 2017) and sums over the estimator for each monomial (Eq. G2). The associated computer code is available through the Github repository.

### Finite-sample estimator for 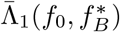

A similar approach be used to obtain an estimator for the “single-rare” case in Eq. (6). The important difference between 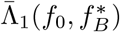 and 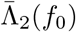 is that we now need to condition the frequency of the *B* allele. We can achieve this by considering a modified version of our previous moment estimator, (Eq. G1) to be

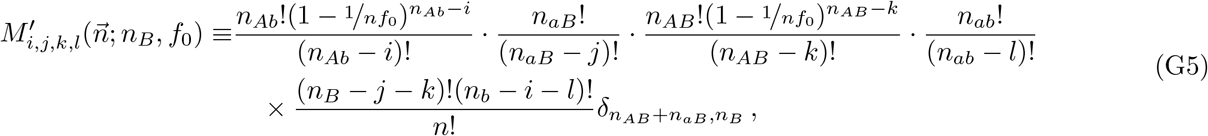

Where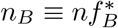, *n*_*b*_ ≡ *n* − *n*_*B*_, and δ_*n,n*_*′* is the Kronecker delta symbol:

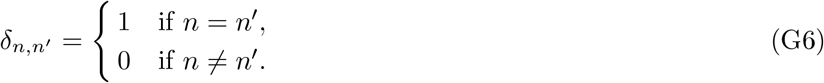

The presence of this delta function constrains the number of *B* alleles to be exactly equal to 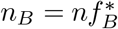. We will show that this modified estimator, when averaged over sampling noise and the stochasticity of the evolutionary dynamics, gives the appropriate frequency-weighted moments,

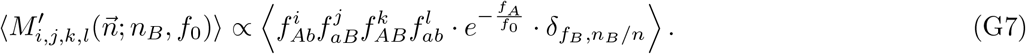

To see this, we first average *M* ′ over the multinomial sampling process, assuming that the haplotype frequencies 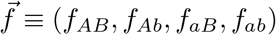 are all held fixed,

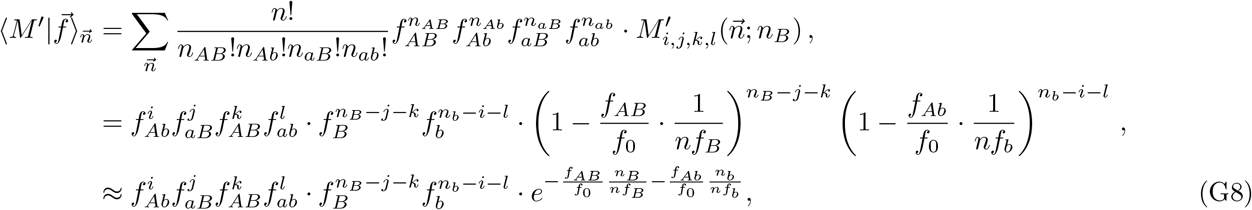

where in the last line we have assumed *n*_*b*_, *n*_*B*_ ≫ *i, j, k, l*. This is justified since we are interested in common *B* alleles, *n*_*B*_ ≫ 1. The term 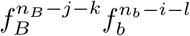 as a function of *f*_*B*_ will be sharply peaked around *f*_*B*_ = *n*_*B*_*/n*, approaching the delta function 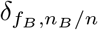 when *n*→ ∞. To see this, we recall that the central limit theorem allows us to approximate the binomial sampling probability with a Gaussian,

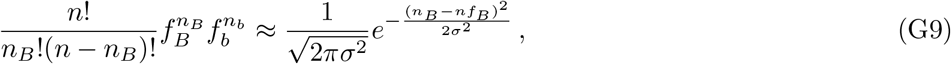

where *σ*^2^ = *nf*_*B*_*f*_*b*_. The Gaussian as a function of *f*_*B*_ approaches the desired delta function because of its vanishing width when *n* → ∞. We therefore obtain the approximation

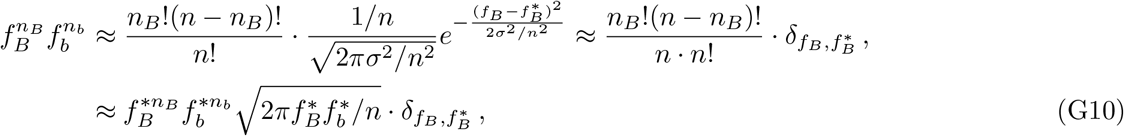

where we have used the Sterling approximation for the factorials.

With the above approximation, we can now average Eq. (G8) over the distribution of haplotype frequencies to obtain

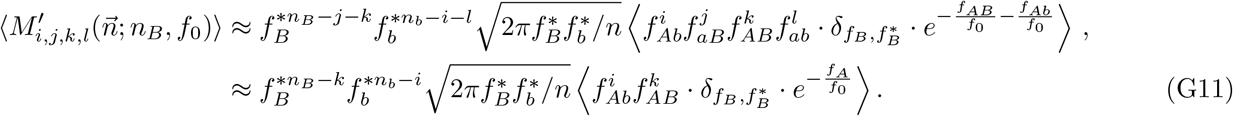

where the last line assumed that 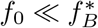, so that *f*_*aB*_ ≈*f*_*B*_ and *f*_*ab*_ ≈*f*_*b*_. As a result, *j* and *l* will not influence the average of *M* ′ in this limit. Using the above identity, we can finally obtain an estimator for 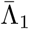

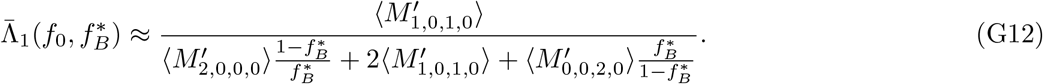

### Finite-sample estimator for *P*(Λ > 0)

In addition to moments like 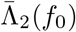 and 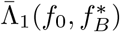, we can also develop finite-sample estimators for other properties of the distribution of Λ, e.g. the probability that all four haplotypes are present (i.e. Λ > 0) in a sample of size *n*, conditioned on observing both alleles near frequency *f*_0_. For simplicity, we will restrict our attention to the recombination-dominated regime in Eq. (18), where the double mutant haplotype is always asymptotically smaller than *f*_0_. When *nf*_0_ ≫ 1, the probability of observing all four haplotypes is therefore equivalent to the probability of observing at least one copy of the double mutant:

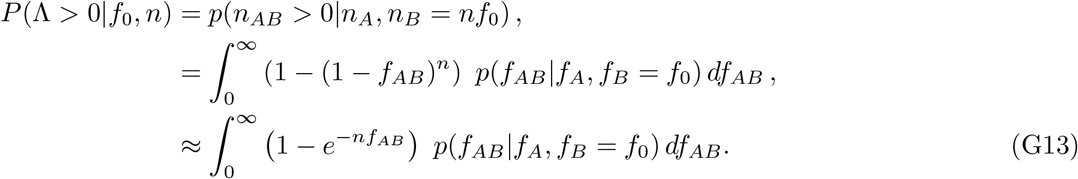

After plugging in the conditional distribution in Eq. (18) and evaluating the integral over *f*_*AB*_, we obtain Eq. (20) in the main text:

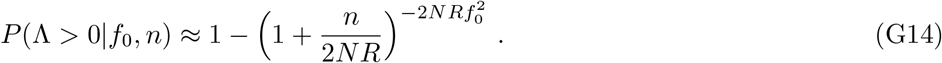

## Appendix H Applications to polymorphism data in E. rectale

To estimate frequency-resolved homoplasy in the commensal human gut bacterium *E. rectale*, we downloaded the genome alignments of a total of 4, 872 non-redundant metagenomically assembled genomes (MAGs) from the Unified Human Gastrointestinal Genome collection (Almeida *et al*., 2021). We focused on protein coding genes that are shared by more than 90% of the genomes in the dataset. Regions shared between overlapping genes were removed. Since our theoretical analysis considered biallelic loci, we filtered out polymorphic sites with more than two alleles. We identified the *A* and *B* alleles to be the minor alleles in the population. We further restricted our attention to synonymous polymorphisms. We recorded the two-site haplotype counts, *n*_*ab*_, *n*_*Ab*_, *n*_*aB*_, *n*_*AB*_, and coordinate distances on the reference genome for all pairs of sites within each gene. We reasoned that at larger coordinate distances, the gene synteny of individual strains might start to differ from the *E. rectale* reference genome, which could make it harder to identify the precise distance between sites.

To include pairs of sites with larger recombination rates, we recorded analogous haplotype counts between 10^3^ pairs of randomly selected genes. These randomly sampled pair of genes are typically separated by hundreds of kilobases in the reference genome, which is longer than the typical recombination length in bacteria (Liu and Good, 2024). This suggests that their recombination rate should approach a constant value, *R*_max_ = lim_*𝓁→∞*_ *R*(𝓁).

## REFERENCES

Almeida, A., S. Nayfach, F. S. Miguel Boland, M. Beracochea, Z. J. Shi et al., 2021 A unified catalog of 204,938 reference genomes from the human gut microbiome. Nature Biotechnology 39: 105–114.

Birzu, G., H. S. Muralidharan, D. Goudeau, R. R. Malmstrom, D. S. Fisher et al., 2023 Hybridization breaks species barriers in long-term coevolution of a cyanobacterial population. eLife 12.

Chan, A. H., P. A. Jenkins, and Y. S. Song, 2012 Genome-wide fine-scale recombination rate variation in Drosophila melanogaster. PLOS Genetics 8: e1003090.

Coop, G., X. Wen, C. Ober, J. K. Pritchard, and M. Przeworski, 2008 High-resolution mapping of crossovers re-veals extensive variation in fine-scale recombination patterns among humans. Science 319: 1395–1398.

Corbett-Detig, R. B., J. Zhou, A. G. Clark, D. L. Hartl, and J. F. Ayroles, 2013 Genetic incompatibilities are widespread within species. Nature 504: 135–137.

Didelot, X., and D. Falush, 2007 Inference of bacterial microevolution using multilocus sequence data. Genetics 175: 1251–1266.

Didelot, X., and D. J. Wilson, 2015 ClonalFrameML: Efficient inference of recombination in whole bacterial genomes. PLOS Computational Biology 11: e1004041.

Eberle, M. A., M. J. Rieder, L. Kruglyak, and D. A. Nickerson, 2006 Allele frequency matching between SNPs reveals an excess of linkage disequilibrium in genic regions of the human genome. PLOS Genetics 2: e142.

Ewens, W. J., 2004 Mathematical Population Genetics: Theoretical Introduction, volume 1. Springer.

Falush, D., T. Wirth, B. Linz, J. K. Pritchard, M. Stephens et al., 2003 Traces of human migrations in Helicobacter pylori populations. Science 299: 1582–1585.

Fay, J. C., and C.-I. Wu, 2000 Hitchhiking under positive Darwinian selection. Genetics 155: 1405–1413.

Fisher, D. S., 2007 Course 11 Evolutionary dynamics. In J.-P. Bouchaud, M. Mézard and J. Dalibard, editors, Les Houches, volume 85 of Complex Systems. Elsevier, 395–446.

Fu, Y. X., and W. H. Li, 1993 Statistical tests of neutrality of mutations. Genetics 133: 693–709.

Garcia, J. A., and K. E. Lohmueller, 2021 Negative linkage disequilibrium between amino acid changing variants reveals interference among deleterious mutations in the human genome. PLOS Genetics 17: e1009676.

Garud, N. R., B. H. Good, O. Hallatschek, and K. S. Pollard, 2019 Evolutionary dynamics of bacteria in the gut microbiome within and across hosts. PLOS Biology 17: e3000102.

Garud, N. R., P. W. Messer, E. O. Buzbas, and D. A. Petrov, 2015 Recent selective sweeps in North American Drosophila melanogaster show signatures of soft sweeps. PLOS Genetics 11: e1005004.

Garud, N. R., P. W. Messer, and D. A. Petrov, 2021 Detection of hard and soft selective sweeps from Drosophila melanogaster population genomic data. PLOS Genetics 17: e1009373.

Good, B. H., 2022 Linkage disequilibrium between rare mutations. Genetics 220: iyac004.

Gutenkunst, R. N., R. D. Hernandez, S. H. Williamson, and C. D. Bustamante, 2009 Inferring the joint demographic history of multiple populations from multidimensional SNP frequency data. PLOS Genetics 5: e1000695.

Halldorsson, B. V., H. P. Eggertsson, K. H. S. Moore, H. Hauswedell, O. Eiriksson et al., 2022 The sequences of 150,119 genomes in the UK Biobank. Nature 607: 732–740.

Harris, R. B., A. Sackman, and J. D. Jensen, 2018 On the unfounded enthusiasm for soft selective sweeps II: examining recent evidence from humans, flies, and viruses. PLOS Genetics 14: e1007859.

Hermisson, J., and P. S. Pennings, 2017 Soft sweeps and beyond: understanding the patterns and probabilities of selection footprints under rapid adaptation. Methods in Ecology and Evolution 8: 700–716.

Hey, J., and J. Wakeley, 1997 A coalescent estimator of the population recombination rate. Genetics 145: 833–846.

Hill, W. G., and A. Robertson, 1968 Linkage disequilibrium in finite populations. TAG. Theoretical and applied genetics. Theoretische und angewandte Genetik 38: 226–231.

Hudson, R. R., 2001 Two-locus sampling distributions and their application. Genetics 159: 1805–1817.

Hudson, R. R., and N. L. Kaplan, 1985 Statistical properties of the number of recombination events in the history of a sample of DNA sequences. Genetics 111: 147–164.

Kim, B. Y., C. D. Huber, and K. E. Lohmueller, 2017 Inference of the distribution of selection coefficients for new nonsynonymous mutations using large samples. Genetics 206: 345–361.

Lewontin, R. C., 1964 The interaction of selection and linkage. I. General considerations. Genetics 49: 49–67.

Li, H., and R. Durbin, 2011 Inference of human population history from individual whole-genome sequences. Nature 475: 493–496.

Lin, M., and E. Kussell, 2017 Correlated mutations and homologous recombination within bacterial populations. Genetics 205: 891–917.

Liu, Z., and B. H. Good, 2024 Dynamics of bacterial recombination in the human gut microbiome. PLOS Biology 22: e3002472.

Lynch, M., S. Xu, T. Maruki, X. Jiang, P. Pfaffelhuber et al., 2014 Genome-wide linkage-disequilibrium profiles from single individuals. Genetics 198: 269–281.

Lynch, M., Z. Ye, L. Urban, T. Maruki, and W. Wei, 2022 The linkage-disequilibrium and recombinational landscape in Daphnia pulex. Genome Biology and Evolution 14: evac145.

Mah, J. C., K. E. Lohmueller, and N. Garud, 2023 Inference of the demographic histories and selective effects of human gut commensal microbiota over the course of human history.

McVean, G. A. T., 2002 A genealogical interpretation of linkage disequilibrium. Genetics 162: 987–991.

McVean, G. A. T., S. R. Myers, S. Hunt, P. Deloukas, D. R. Bentley et al., 2004 The fine-scale structure of recombination rate variation in the human genome. Science 304: 581–584.

Meurer, A., C. P. Smith, M. Paprocki, O. Čertík, S. B. Kirpichev et al., 2017 Sympy: Symbolic computing in python. PeerJ Computer Science 3: e103.

Myers, S., L. Bottolo, C. Freeman, G. McVean, and P. Donnelly, 2005 A fine-scale map of recombination rates and hotspots across the human genome. Science 310: 321–324.

Myers, S. R., and R. C. Griffiths, 2003 Bounds on the minimum number of recombination events in a sample history. Genetics 163: 375–394.

Neher, R. A., and T. Leitner, 2010 Recombination rate and selection strength in HIV intra-patient evolution. PLOS Computational Biology 6: 1–7.

Ohta, T., and M. Kimura, 1971 Linkage disequilibrium between two segregating nucleotide sites under the steady flux of mutations in a finite population. Genetics 68: 571–580.

Ragsdale, A. P., 2022 Local fitness and epistatic effects lead to distinct patterns of linkage disequilibrium in protein-coding genes. Genetics 221: iyac097.

Ragsdale, A. P., and S. Gravel, 2019 Models of archaic admixture and recent history from two-locus statistics. PLOS Genetics 15: e1008204.

Ragsdale, A. P., C. Moreau, and S. Gravel, 2018 Genomic inference using diffusion models and the allele frequency spectrum. Current Opinion in Genetics and Development 50: 140–147.

Ragsdale, A. P., T. D. Weaver, E. G. Atkinson, E. G. Hoal, M. Möller et al., 2023 A weakly structured stem for human origins in Africa. Nature 617: 755–763.

Romero, E. V., and A. F. Feder, 2024 Elevated HIV viral load is associated with higher recombination rate in vivo. Molecular Biology and Evolution 41: msad260.

Rosen, M. J., M. Davison, D. Bhaya, and D. S. Fisher, 2015 Fine-scale diversity and extensive recombination in a quasisexual bacterial population occupying a broad niche. Science 348: 977–978.

Sabeti, P. C., D. E. Reich, J. M. Higgins, H. Z. P. Levine, D. J. Richter et al., 2002 Detecting recent positive selection in the human genome from haplotype structure. Nature 419: 832–837.

Santiago, E., I. Novo, A. F. Pardiñas, M. Saura, J. Wang et al., 2020 Recent demographic history inferred by high-resolution analysis of linkage disequilibrium. Molecular Biology and Evolution 37: 3642–3653.

Sawyer, S. A., and D. L. Hartl, 1992 Population genetics of polymorphism and divergence. Genetics 132: 1161–1176.

Slatkin, M., 2008 Linkage disequilibrium — understanding the evolutionary past and mapping the medical future. Nature Reviews Genetics 9: 477–485.

Sohail, M., O. A. Vakhrusheva, J. H. Sul, S. L. Pulit, L. C. Francioli et al., 2017 Negative selection in humans and fruit flies involves synergistic epistasis. Science 356: 539–542.

Song, Y. S., and J. S. Song, 2007 Analytic computation of the expectation of the linkage disequilibrium coefficient r2. Theoretical Population Biology 71: 49–60.

Spence, J. P., and Y. S. Song, 2019 Inference and analysis of population-specific fine-scale recombination maps across 26 diverse human populations. Science Advances 5: eaaw9206.

Stephan, W., Y. S. Song, and C. H. Langley, 2006 The hitchhiking effect on linkage disequilibrium between linked neutral loci. Genetics 172: 2647–2663.

Sun, K. Y., X. Bai, S. Chen, S. Bao, M. Kapoor et al., 2023 A deep catalog of protein-coding variation in 985,830 individuals.

Suzuki, T. A., J. L. Fitzstevens, V. T. Schmidt, H. Enav, K. E. Huus et al., 2022 Codiversification of gut microbiota with humans. Science 377: 1328–1332.

Tajima, F., 1989 Statistical method for testing the neutral mutation hypothesis by DNA polymorphism. Genetics 123: 585–595.

Turakhia, Y., B. Thornlow, A. Hinrichs, J. McBroome, N. Ayala et al., 2022 Pandemic-scale phylogenomics reveals the SARS-CoV-2 recombination landscape. Nature 609: 994–997.

Vakhrusheva, O. A., E. A. Mnatsakanova, Y. R. Galimov, T. V. Neretina, E. S. Gerasimov et al., 2020 Genomic signatures of recombination in a natural population of the bdelloid rotifer Adineta vaga. Nature Communications 11: 6421.

Virtanen, P., R. Gommers, T. E. Oliphant, M. Haberland, T. Reddy et al., 2020 SciPy 1.0: Fundamental algorithms for scientific computing in Python. Nature Methods 17: 261–272.

Wakeley, J., 2008 Conditional gene genealogies under strong purifying selection. Molecular Biology and Evolution 25: 2615–2626.

Walczak, A. M., L. E. Nicolaisen, J. B. Plotkin, and M. M. Desai, 2012 The structure of genealogies in the presence of purifying selection: A fitness-class coalescent. Genetics 190: 753–779.

Weissman, D. B., M. M. Desai, D. S. Fisher, and M. W. Feldman, 2009 The rate at which asexual populations cross fitness valleys. Theoretical Population Biology 75: 286–300.

Weissman, D. B., M. W. Feldman, and D. S. Fisher, 2010 The rate of fitness-valley crossing in sexual populations. Genetics 186: 1389–1410.

Wolff, R., and N. R. Garud, 2023 Pervasive selective sweeps across human gut microbiomes.

Zanini, F., J. Brodin, L. Thebo, C. Lanz, G. Bratt et al., 2015 Population genomics of intrapatient HIV-1 evolution. eLife 4: e11282.

